# AntiBP3: A hybrid method for predicting antibacterial peptides against gram-positive/negative/variable bacteria

**DOI:** 10.1101/2023.07.25.550443

**Authors:** Nisha Bajiya, Shubham Choudhury, Anjali Dhall, Gajendra P. S. Raghava

## Abstract

This study focuses on the development of in silico models for predicting antibacterial peptides as a potential solution for combating antibiotic-resistant strains of bacteria. Existing methods for predicting antibacterial peptides are mostly designed to target either gram-positive or gram-negative bacteria. In this study, we introduce a novel approach that enables the prediction of antibacterial peptides against several bacterial groups, including gram-positive, gram-negative, and gram-variable bacteria. Firstly, we developed an alignment-based approach using BLAST to identify antibacterial peptides and achieved poor sensitivity. Secondly, we employed a motif-based approach to predict antibacterial peptides and obtained high precision with low sensitivity. To address the similarity issue, we developed machine learning-based models using a variety of compositional and binary features. Our machine learning-based model developed using the amino acid binary profile of terminal residues achieved maximum AUC 0.93, 0.98 and 0.94 for gram-positive, gram-negative, and gram-variable bacteria, respectively, when evaluated on a validation/independent dataset. Our attempts to develop hybrid or ensemble methods by merging machine learning models with similarity and motif-based techniques did not yield any improvements. To ensure robust evaluation, we employed standard techniques such as five-fold cross-validation, internal validation, and external validation. Our method performs better than existing methods when we compare our method with existing approaches on an independent dataset. In summary, this study makes significant contributions to the field of antibacterial peptide prediction by providing a comprehensive set of methods tailored to different bacterial groups. As part of our contribution, we have developed the AntiBP3 web server and standalone package, which will assist researchers in the discovery of novel antibacterial peptides for combating bacterial infections (https://webs.iiitd.edu.in/raghava/antibp3/).

**Key Points:**

⍰ BLAST-based similarity for annotating antibacterial peptides.
⍰ Machine learning-based models developed using composition and binary profiles.
⍰ Identification and mapping of motifs exclusively found in antibacterial peptides
⍰ Improved version of AntiBP and AntiBP2 for predicting antibacterial peptides.
⍰ Web server for predicting/designing/scanning antibacterial peptides for all groups of bacteria

**Author’s Biography:**

1. Nisha Bajiya is currently working as Ph.D. in Computational Biology from Department of Computational Biology, Indraprastha Institute of Information Technology, New Delhi, India.
2. Shubham Choudhury is currently working as Ph.D. in Computational Biology from Department of Computational Biology, Indraprastha Institute of Information Technology, New Delhi, India.
3. Anjali Dhall is currently working as Ph.D. in Computational Biology from Department of Computational Biology, Indraprastha Institute of Information Technology, New Delhi, India.
4. Gajendra P. S. Raghava is currently working as Professor and Head of Department of Computational Biology, Indraprastha Institute of Information Technology, New Delhi, India.

## Introduction

Since their initial discovery, the widespread use of antibiotics has resulted in the emergence of drug-resistant strains of pathogenic bacteria [1]. This has led to the development of “superbugs” resistant to existing antibiotics. Moreover, the process of identifying novel compounds for drug design requires substantial financial and temporal investments. Additionally, the growing resistance observed in pathogenic strains, coupled with the side effects associated with antibiotics, highlights the need for alternative treatment options that are both less toxic and less likely to induce resistance. Peptide-based therapeutics have gained considerable attention as one such alternative. Antimicrobial peptides (AMPs) are a class of small oligopeptides that exhibit diverse protective activities against pathogenic microbes [2, 3]. These peptides are categorized based on their specific activities, including antibacterial, antiviral, antiparasitic, and antifungal properties [4].

Antibacterial peptides (ABPs) are short oligopeptides of between 5 and 100 amino acid residues. They are distinguished by their cationic nature and enriched with certain amino acids such as Arginine and Lysine. These peptides have amphipathic properties, which aid in their incorporation into pathogen cell membranes. (as illustrated in Figure 1) [5]. ABPs can be classified into two categories based on their synthesis types: ribosomally or non-ribosomally synthesized. They display a wide range of structural diversity, including β-sheets, random coils, extended structures, cyclic peptides, and α-helices. Some ABPs also contain disulfide bridges [6]. The antibacterial mode of action exhibited by these peptides involves direct killing through membrane damage. This leads to the formation of patches on the microbial membranes, resulting in cytoplasmic leakage, osmotic shock, and, ultimately, the death of the pathogen. Additionally, ABPs can employ intracellular modes of action, such as modulating enzymatic activity, protein degradation & synthesis, and nucleic acid synthesis. This multifaceted mode of action provides an advantage over traditional antibiotics in terms of reduced development of resistance [2, 4, 7, 8].

**Figure 1:**
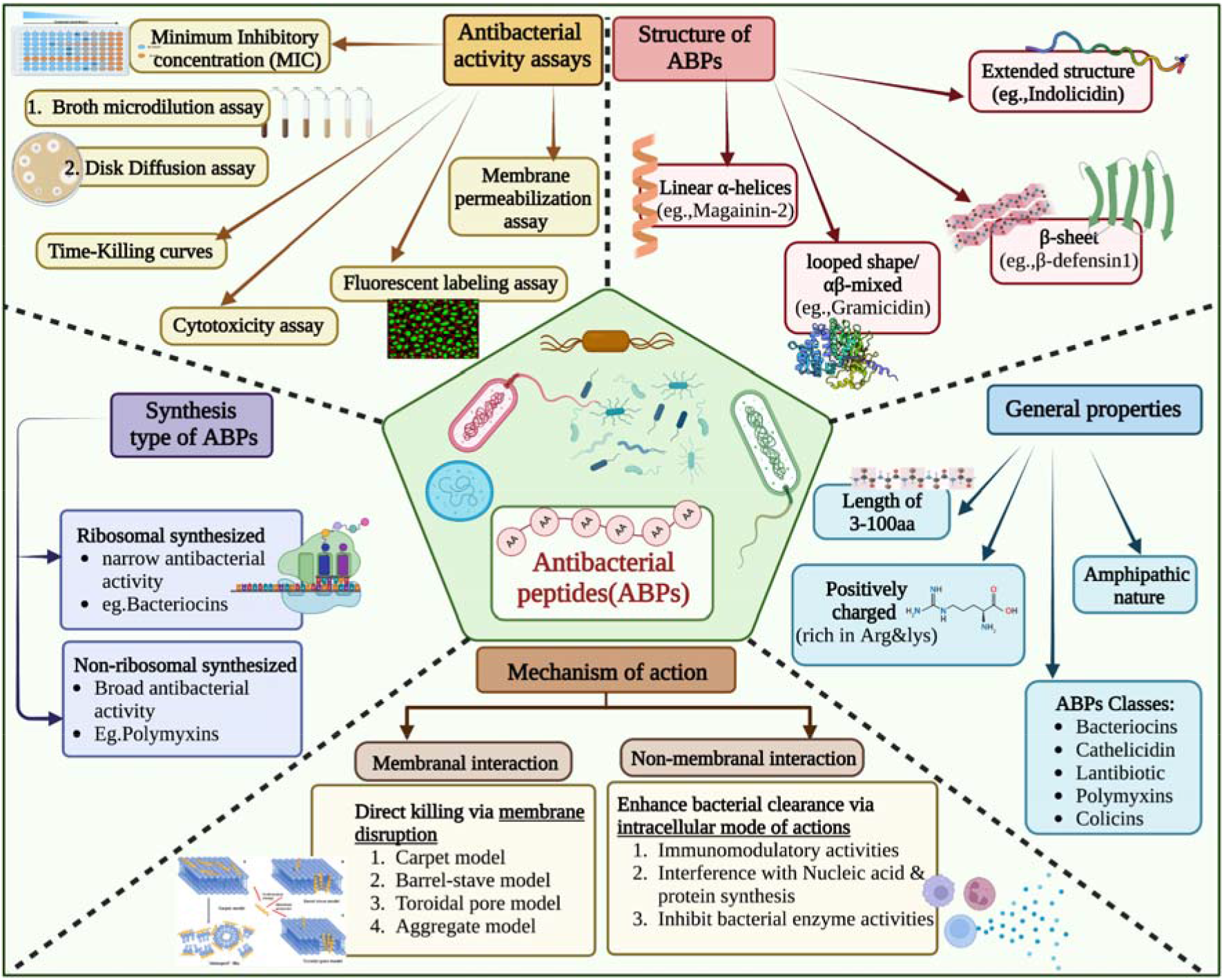
Shows overview of Antibacterial Peptides: Properties, Mechanisms, Assays, Structures, and Synthesis Type.

In the last few decades, considerable progress has been made in the field of ABPs, and experimental methods, as depicted in Figure 1, remain the gold standard for determining the antibacterial activity of peptides. However, these methods are time-consuming, labour-intensive, and expensive, presenting numerous challenges [9, 10,11]. Therefore, in the initial stages of drug design, screening of millions of peptides is crucial to facilitate the discovery of novel ABPs [12]. In order to overcome this challenge, numerous in silico methods have been developed for predicting antibacterial peptides. In 2007, Lata et al. developed a method of AntiBP using machine learning (ML) techniques to classify antibacterial and non-antibacterial peptides, where they employed Support Vector Machine (SVM), Quantitative Matrix (QM) and Artificial Neural Network (ANN) and [13]. Since then, numerous prediction models have been developed that include AmpGram [14], AI4AMP [15], AMPfun[16], AMPScanner[17, 18], Deep-AmPEP30[19], iAMPpred [20], ACEP [21], Deep-ABPpred [22], and StaBle-ABPpred [23]. In addition, attempts have been made to develop class-specific prediction servers like AntiBP2 for predicting the source of antibacterial peptides [24]. AMPScanner vr.2 [18] allows the identification of AMPs active against gram-positive and/or gram-negative bacteria. YB Ruiz-Blanco et al. developed ABP-Finder, a two-stage classifier that classifies peptides as ABPs and further distinguishes them as either gram-positive or gram-negative peptides [12]. Söylemez, Ü. G. et al. proposed an RF-based method that utilizes ten different physicochemical properties and 100-fold Monte Carlo Cross-Validation to classify linear cationic AMPs into gram-negative and gram-positive peptides [25, 26].

One of the most significant disadvantages of present approaches is their inability to address all bacterial types, notably gram-(indeterminate/variable) bacteria, which cannot be detected by gram-staining methods, leaving a gap in the analysis of antibacterial peptides. Furthermore, the models are often trained on limited datasets that are no longer a representation of the current state of the field. In order to complement the existing method, a conscious and systematic effort was undertaken to address the aforementioned shortcomings. Our method is inspired by the ABP-Finder tool, where they have covered peptides with a gram-positive, gram-negative, and broad spectrum of peptides having properties of both; it excludes gram-variable bacterial species. Therefore, our research broadens the scope by introducing a third group, gram-variable antibacterial peptides (GV ABPs), which contain ABPs against gram-variable bacterial species as well as peptides with a broad spectrum of properties. The focus of this study is to build prediction models capable of identifying antibacterial peptides for all categories of bacteria, including gram-variable strains, by leveraging large and diverse datasets. As a result, we created three separate datasets from public databases, each of which contained balanced and high-quality experimentally validated antibacterial peptides. Classification models were created for each bacterial group, and their performance was measured using various metrics such as accuracy, precision, specificity, sensitivity, recall, Area Under Curve (AUC), F1 measure, and Matthews correlation coefficient (MCC).

## MATERIAL AND METHODS

### 1. Dataset

We collected the positive dataset (i.e., antibacterial peptides) from multiple repositories that specialize in antibacterial peptides for gram-positive (GP), gram-negative (GN), and gram-variable (GV) bacteria. These public repositories include the Antimicrobial Peptide Database (APD)[27], AntiBP2 [24], dbAMP 2.0 [28], Collection of Anti-Microbial Peptides (CAMPR3) [29], Data Repository of Antimicrobial Peptides (DRAMP) [30] and ABP-Finder [12], as depicted in Figure 2. To ensure uniqueness and consistency, we removed similar sequences found in different databases. Our focus was on sequences containing standard amino acids, so any sequences containing non-standard amino acids (BJOUXZ) were also excluded. In vitro synthesis of long peptides is challenging, and larger ABPs tend to exhibit increased toxicity and decreased stability [31]. Similarly, very short peptide sequences often lack potent antibacterial activity. Consequently, we eliminated sequences shorter than eight and longer than 50 amino acids. Following the pre-processing steps, our dataset consisted of 11,370 unique antibacterial peptides; belonging to three groups: 930 peptides for GP bacteria, 1455 peptides for GN bacteria, and 8,985 peptides for GV bacteria.

**Figure 2:**
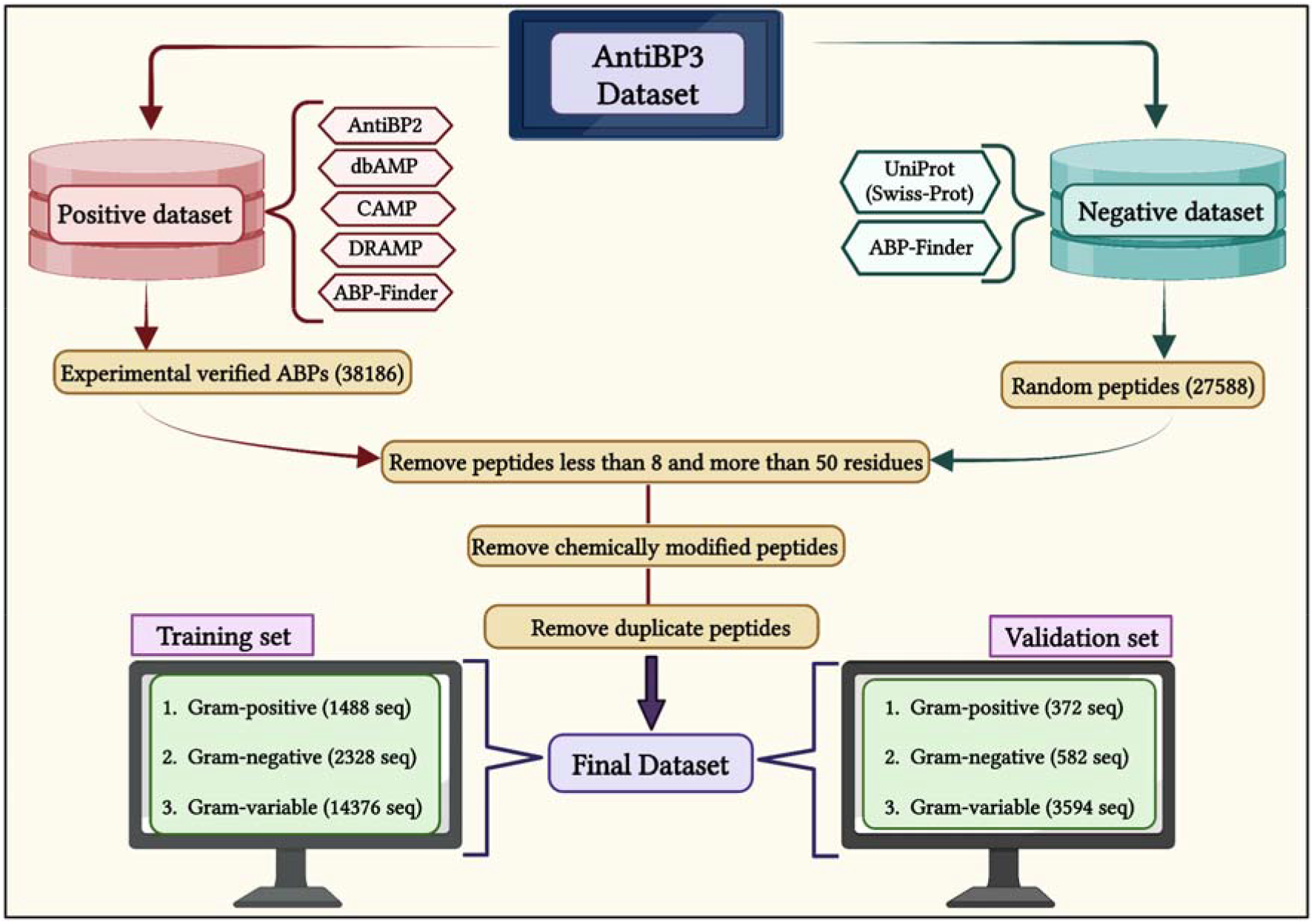
A flow diagram shows the process of creating AntiBP3 datasets for gram-positive, gram-negative, and gram-variable bacteria.

To create non-antibacterial peptides (non-ABPs) for the negative dataset, we followed a similar approach as used in the development of a negative set for AntiBP[13] and AntiBP2 [24]. Since currently, there is no public repository specifically dedicated to non-ABPs; thus, we generate non-antibacterial peptides from the UniProt database [32]. We queried the UniProt database for proteins that met the following criteria: manually reviewed and annotated, consisting of standard amino acid sequences within the range of eight to 50 and lacking any mention of specific keywords associated with antibacterial activity. These keywords included terms such as antibacterial, antitoxin, antimalarial, antitumor, anti-HIV, antiviral, antibiotic antimicrobial, antifungal, antidiabetic, insecticidal, antioxidant, chemotactic, anti-Gram positive, antiprotozoal, anti-Gram negative, antiparasitic, anti-TB, antibiofilm, anti-protist, anti-MRSA, antiparasitic anti-endotoxin, anticancer, anti-inflammatory, antiparasitic, spermicidal, secreted proteins, toxin, excreted, effector, grammistin, endotoxin, bacteriocin, defensin, and lysozyme. This search yielded 11,257 sequences, which were then integrated with the negative dataset used in the ABP-Finder. We then removed the duplicate sequences, resulting in a final set of 10,000 unique sequences called non-ABPs. This non-ABP data then be randomly split according to different classes of ABP’s training and validation set in order to create a balanced dataset.

To enable an unbiased evaluation, we performed the evaluation of models using internal and external cross-validation. We divide the entire data into training and validation datasets, where the training dataset contains 80% of the data, and the validation dataset contains 20% of the data, as shown in Figure 2. In the case of GP, the training dataset comprises 744 ABP and 744 non-ABP, whereas the validation dataset has 186 ABP and 186 non-ABP, resulting in an 80:20 ratio. Similarly, the training dataset for GN contains 1,164 ABP and 1,164 non-ABP; and the validation dataset includes 291 ABP and 291 non-ABP. Likewise, the training dataset for GV contains 7,188 ABP and 7,188 non-ABP, whereas the validation dataset contains 1,797 ABP and 1,797 non-ABP. Therefore, we obtained a balanced dataset with a total final count of 930 ABP & 930 non-ABP for GP bacteria, 1,455 ABP & 1,455 non-ABP for GN bacteria, and 8,985 ABP & 8,985 non-ABP for GV bacteria. The validation dataset was not utilized either for training or testing the method. These validation peptides are provided as an independent dataset for assessing the performance of the prediction models

### 2. Overall workflow

The complete workflow of the current study is illustrated in Figure 3.

**Figure 3:**
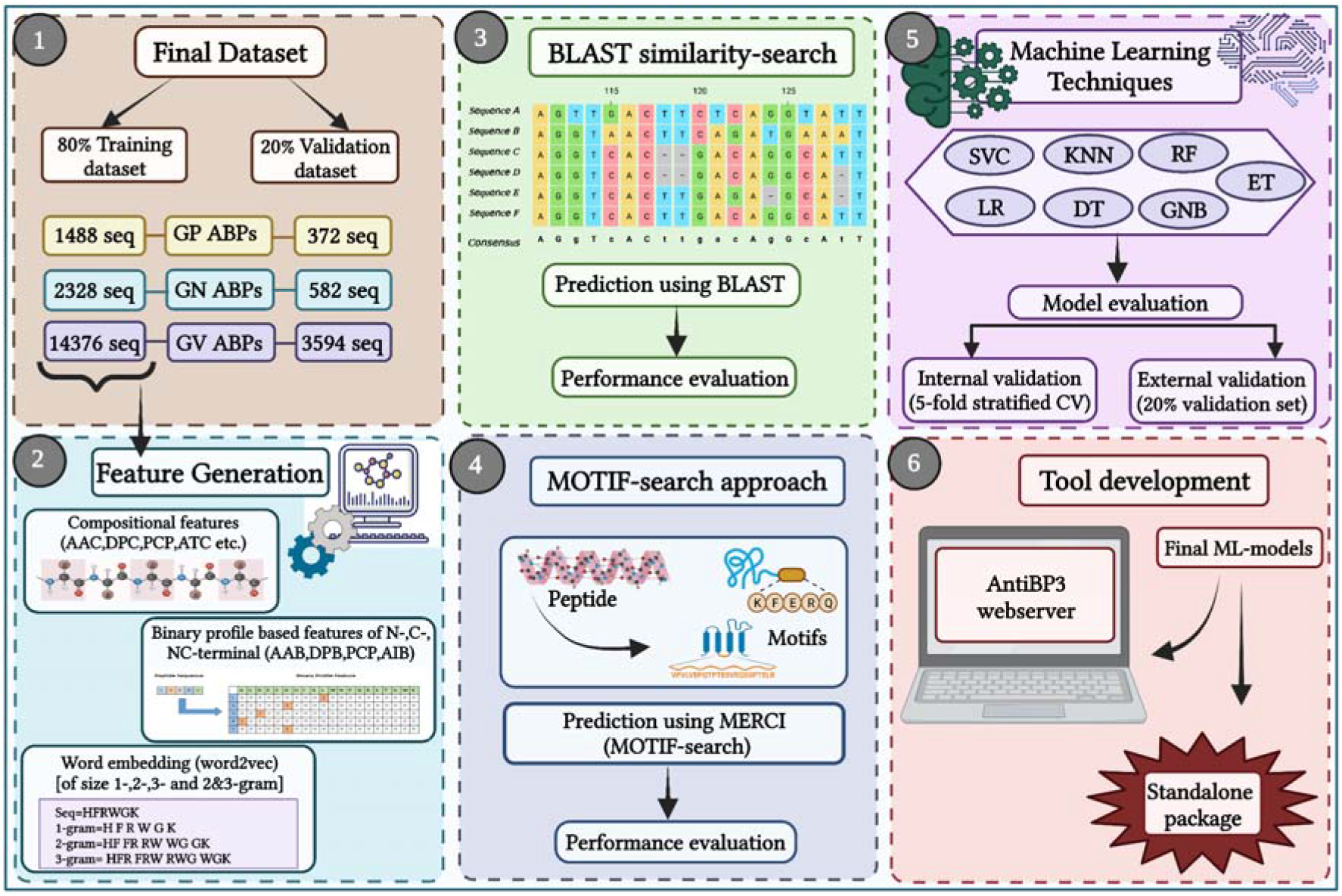
Overall study architecture from feature generation to web server development.

### 3. Compositional analysis of amino acid residues

In the past, general antimicrobial peptide prediction methods/tools relied on different peptide sequence features, such as peptide sequence length and amino-acid level-based features [22]. We used the most relevant characteristics, such as amino acid composition. We have employed Pfeature [33] to compute the amino acid composition (AAC) of training and validation datasets. We built a feature vector of length 20 using the following equation 1, which specifies the percentage composition of 20 amino-acid residues.

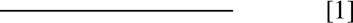

where AAC_i_ and AAR_i_ are the percentage composition of amino acid and the number of specific residues of type *i* in a peptide, respectively.

### 4. Two Sample Logo

We used the Two Sample Logo (TSL) [34] to construct sequence logos for the three categories of ABPs in both the positive and negative datasets. The sequence logos graphically illustrate the relevance of the amino acid residues at each location in the peptide sequence. The x-axis depicts the amino acid residues in the generated sequence logos. At the same time, the positive y-axis displays the bit-score for the enriched residues, whereas the negative y-axis displays the depleted residues in the given peptide sequences, reflecting the relevance of a single residue at a given location. TSL, on the other hand, accepts peptide sequences as a fixed-length vector for input. Considering the shortest peptide length in our dataset, which is eight residues, we only account for eight amino acids from the N-terminus (beginning) and eight residues from the C-terminus (end) for each peptide sequence. We combined these two segments to create a fixed-length vector of sixteen residues. This fixed-length vector format was used in our study on all three different groups of ABPs to create sequence logos.

### 5. Sequence alignment method

In this study, we used an alignment-based approach to categorise ABPs into GP, GN, and GV based on similarity. We utilized the NCBI-BLAST+ version 2.13.0+ (blastp suite) to implement a similarity search approach [35, 36]. First, we used the makeblastdb suite of NCBI-BLAST+ to construct three custom databases. We created a specialized ABP database containing the respective ABP sequences and a non-ABPs database containing non-ABP sequences for each group set (GP, GN, and GV). Finally, the validation dataset was then queried against the custom databases using the BLASTP suite. The peptides were classified based on the presence of the most significant hit in either the respective ABPs or the non-ABPs databases. If the top BLAST result was an ABPs peptide from a certain group, the query sequence was classified as a member of that group. On the other hand, if the top hit was from the non-ABPs database, the peptide was classified as a non-ABPs sequence. We conducted the BLAST with several e-value cut-offs ranging from 1e-20 to 1e+3 to discover the optimal value of the e-value threshold. We intended to find the optimal e-value cut-off that produced the greatest classification performance for our ABPs prediction job by examining the results at various e-value thresholds.

### 6. Motif search

In our study, we investigated the motif-based strategy for identifying conserved motifs in ABPs. While the motif-based technique has previously been used to identify motifs in proteins [37, 38], it has not been widely used to uncover patterns in antibacterial peptides. Therefore, we employed Motif—EmeRging and with Classes—Identification (MERCI) [39], a tool for identifying conserved motifs. We separated the dataset into positive and negative subsets for each ABP group to conduct the analysis. In MERCI, both the positive and negative datasets were given as inputs but only provided motifs for the positive instances. The algorithm then found ungapped patterns/motifs with a motif occurrence frequency of 10, 20, and 30 (fp10, fp20, and fp30) that may successfully differentiate between positive and negative samples for the particular ABP group. Thus, for each ABP group, we obtained three distinct sets of motifs and computed the overall motif coverage. The list of unique motifs for each group of ABP is provided on our web server.

### 7. Feature generation

In the current work, we estimated various features utilizing peptide sequence information. We utilised the Pfeature [33] standalone software to compute composition-based and binary profile-based features of our datasets.

#### 7.1. Compositional features

For each peptide sequence in both positive and negative datasets, a total of 2534 compositional features have been generated. We calculated seventeen different types of descriptors/features, including AAC (Amino acid composition), APAAC (Amphiphilic pseudo amino acid composition), DDR (Distance distribution of residue), DPC (Di-peptide composition), QSO (Quasi-sequence order), PCP (Physico-chemical properties composition), PAAC (Pseudo amino acid composition), RRI (Residue repeat Information), SPC (Shannon entropy of physicochemical properties), ATC (Atomic composition), BTC (Bond type composition), CTC (Conjoint triad descriptors), AAI(Amino Acid index), PRI(Property repeats index), SEP(Shannon entropy of a protein), SER(Shannon entropy of a residue), SOC (Sequence order coupling number)etc. In this study, we developed prediction models using each feature.

#### 7.2. Binary profile features

In order to compute binary profiles of ABPs and non-ABPs, the length of variables should be fixed. The minimum length of peptides in our dataset is 8; hence we constructed binary profiles for N8, C8, and combined terminal residues N8C8 after computing the fixed length patterns. Similarly, for each peptide sequence in both positive and negative datasets, we generated four different types of binary profile features for N-, C-, and NC-terminals, including AAB (Amino acid-based binary profile), DPB (Dipeptide-based binary profile), PCB (Physico-chemical properties based binary profile), AIB (Amino-acid indices based binary profile) etc. We developed prediction methods for each of the binary features mentioned above in our study.

#### 7.3. Word embedding

Word embedding is a strong natural language processing technique that includes encoding words as vectors in a high-dimensional space based on their contextual information [40]. The FastText method was used as our word embedding technique[41, 42]. Using information from a huge collection of text documents, the fastText method provides similar vector representations to words that appear in similar contexts. This enables us to capture semantic relationships as well as word similarities. We carried out preprocessing procedures on the peptide sequences to prepare our dataset for training. Furthermore, the peptides were converted into n-grams of varying sizes, such as 1-grams (individual words), 2-grams (pairs of nearby words), 3-grams (triplets of adjacent words), and a mixture of 2-grams and 3-grams. We next trained the fastText model on the preprocessed training dataset, which generates word embedding vectors with a fixed dimension (often several hundred) [43]. The semantic and contextual information of the words is captured by these vectors. Using these word embedding vectors, we turned our peptide sequences into feature vectors. Each feature vector represents a peptide and includes information about the words and their contextual relationships. Finally, we used these feature vectors to train our model to accomplish the necessary classification or prediction job.

### 8. Machine Learning Algorithms

Several machine learning methods were used to create the three classification models in our study. Random Forest (RF), Decision Tree (DT), Gaussian Naive Bayes (GNB), Logistic Regression (LR), Support Vector Classifier (SVC), K-Nearest Neighbour (kNN), and Extra Tree (ET) are among these methods. To build these classifiers, we used the Scikit-learn package, a prominent Python library for machine learning. By recursively dividing the dataset depending on feature requirements, the DT method creates a tree-like model for classification or regression tasks [44]. RF, on the other hand, are ensemble learning approaches that construct numerous decision trees during the training phase and aggregate their predictions to achieve correct estimations [45].

SVM is a machine-learning approach that uses vector space to identify a decision boundary that maximizes the separation between two classes [46]. The KNN algorithm is a supervised learning technique that is used for classification and regression applications. It identifies the class or value of a new data point based on its closeness to the training dataset’s k closest neighbours [47]. GNB is a probabilistic classification technique that employs Bayes’ theorem with the assumption of strong feature independence [48]. Another ensemble learning approach based on decision trees is the ET classifier. ET Classifier, like RF, randomizes some decisions and data subsets to prevent overfitting [49]. In addition to these algorithms, we also employed LR, a supervised learning algorithm that learns linear functions to predict outcomes [50].

### 9. Cross-validation Techniques

We employed both internal and external validation strategies to evaluate the performance of our models. For internal validation, we used the stratified five-fold cross-validation approach, which helps to reduce biases and overfitting. The training dataset was randomly split into five equal sets, each with a comparable number of ABPs and non-ABPs. Four of the five sets were utilised for training the models, while the fifth set was used for testing. This method was done five times, with each set acting as the test set once, ensuring robust evaluation across multiple iterations [51]. The models’ performance was then assessed using the average performance throughout the five test sets. We used a 20% independent validation dataset that was distinct from the training dataset for external validation. The performance of the best model generated using the training dataset was evaluated using this independent dataset. We got a measure of the model’s generalisation power and ability to reliably classify ABPs by evaluating its performance on unseen data. We aimed to verify the reliability and efficacy of our ABP prediction classification models by using these internal and external validation methodologies.

### 10. Cross-prediction Analysis

In order to understand the need to develop three separate models for GP, GN and GV ABPs, we have done cross-prediction of the models. For example, the model developed for gram-positive ABPs with the best feature and best classifier is used to make predictions on the gram-negative ABPs validation dataset. Similarly, this has been done for all three models developed for GP, GN and GV ABPs to ensure the necessity for three individual models.

### 11. Hybrid or Ensemble Approach

In our study, we also used a hybrid or ensemble technique. The following two hybrid techniques were applied in this case: (i) the alignment-based method (BLAST) was combined with the alignment-free approach (ML-based prediction), and (ii) the motif-based approach (MERCI) was combined with ML-based prediction. In the first strategy, we categorise the peptide using BLAST. Following that, for each right positive prediction, i.e., ABP, we incorporate a ’+0.5’ value and a ’-0.5’ score for the negative predictions, i.e., non-ABP, and a ’0’ score if no hit was identified. The peptide sequence was classified using MERCI in the second method. Similarly, we awarded a score of ’+0.5’ if the motifs were identified and a score of ’0’ if they were not detected. We also incorporated the prediction score produced with machine learning-based algorithms. Finally, we integrate the BLAST and MOTIF scores with the machine learning prediction scores, respectively, to produce final predictions. At varied thresholds, the hybrid approach’s total score was utilised to classify the peptides as antibacterial or non-antibacterial. In the past, several researchers have made substantial use of hybrid methodologies that combine various approaches [36, 52]

### 12. Performance Evaluation

The performance of several models was tested using conventional performance assessment measures. We calculated both threshold-dependent metrics (such as sensitivity, specificity, accuracy, Matthews Correlation Coefficient (MCC), and F1-score) and independent parameters such as Area Under Receiver Operating Characteristics (AUCROC) and Area Under the Precision-Recall Curve (AUPRC). The evaluation parameter formulas are shown in the following equations (2-6).

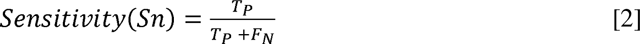

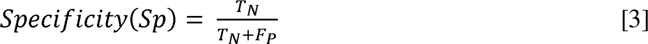

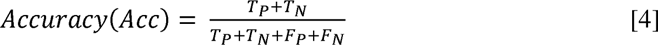

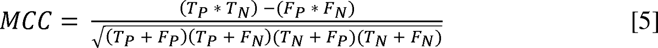

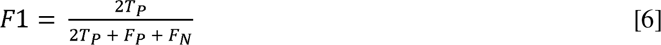

Where TP and TN are successfully predicted ABPs and non-ABPs, while FN is incorrectly predicted ABPs and non-ABPs. Sensitivity (Sn) is the percentage of antibacterial peptides predicted as antibacterial peptides; specificity (Sp) is the percentage of non-antibacterial peptides predicted as non-antibacterial peptides; overall accuracy is the percentage of correctly predicted antibacterial and non-antibacterial peptides. When the positive and negative samples in the data set are out of balance, MCC is used to assess the predictor’s performance. Its value ranges from −1 to 1; a higher MCC indicates a more accurate prediction. The harmonic means of the accuracy and recall scores are used to determine the F1 score. Its value ranges from 0 to 100%, with a higher F1 score indicating a higher-quality classifier [38].

## Results

### 1. Compositional Analysis

We calculated the composition of amino acids for three groups of ABP training datasets. Following that, we computed the average compositions of ABP and non-ABP residues. The graph in Figure 4 depicts a comparison of compositional analyses for the three groups of ABPs. In the case of the GP ABPs dataset, amino acids such as leu, lys, arg, cys, and trp have a higher average composition compared to non-ABPs. Regarding the GN ABPs, the average composition of residues like leu, lys, arg, trp, and pro is higher. Similarly, for the GV ABPs, the amino acids leu, lys, arg, and trp have a higher composition, respectively, compared to non-ABPs. It is important to note that different types of ABPs (e.g., GP, GN, GV) have different compositions for different amino acids; for example, see the composition of residues cys, lys, and arg. These observations indicate that each group of ABPs have different property.

**Figure 4:**
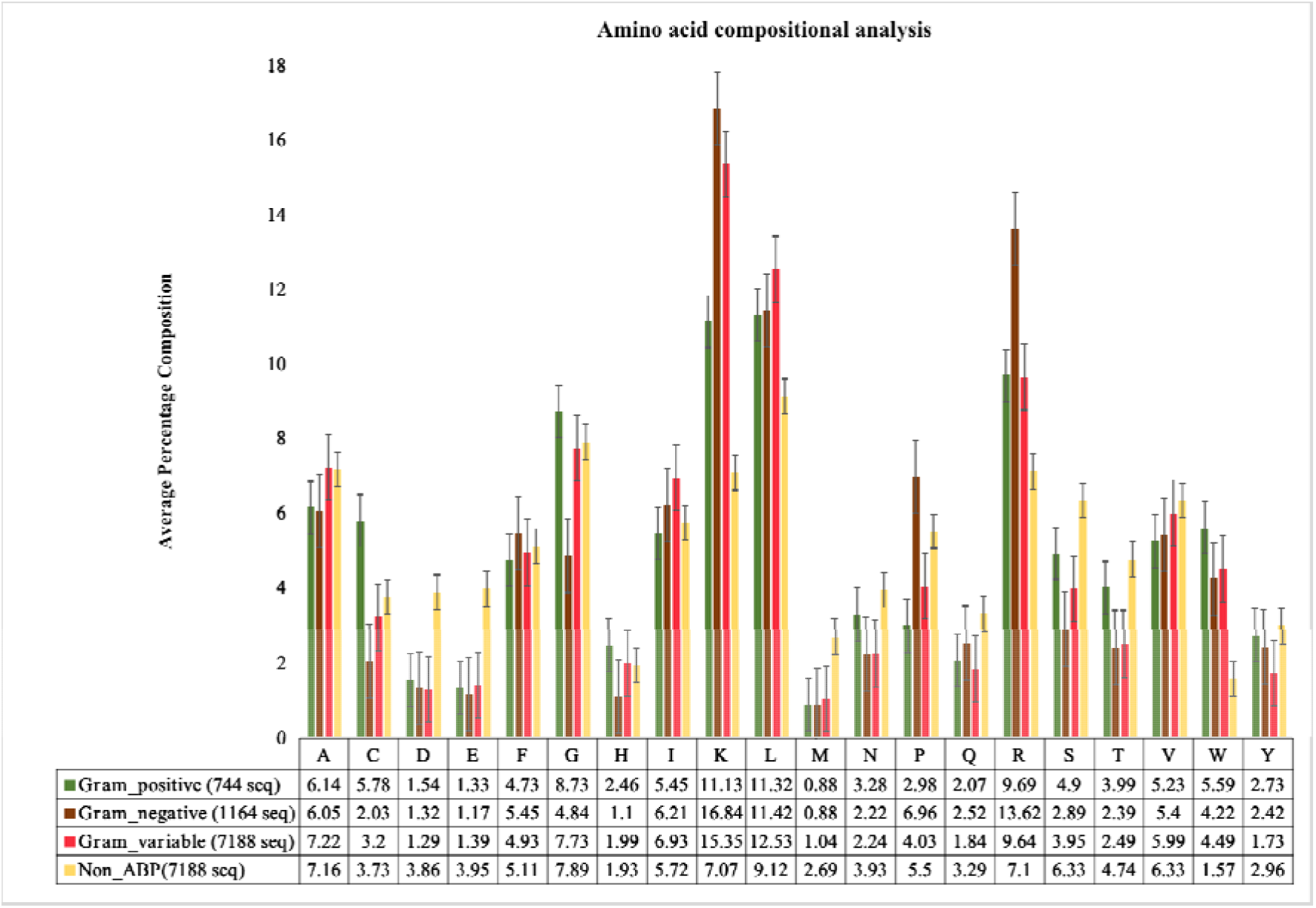
Average amino-acid composition of ABPs and non-ABPs for Gram-positive, negative and variable bacteria.

### 2. Positional Conservation Analysis

Previously, positional studies of antibacterial peptides in AntiBP and AntiBP2 revealed that specific residues in terminal regions were preferred over others. This prompted us to conduct positioning analyses for our three sets of ABPs to determine preference patterns. As depicted in Figure 5, residues like lys, arg, ala, leu, and phe predominate in the first position of GP ABPs at the N-terminus, whereas leu, arg, and trp are commonly found in the second position, while the C-terminus of GP ABPs is dominated by residues gly, trp, arg, and lys. At the N-terminus of GN ABPs, residues like lys, val, arg, leu, and phe predominated in the first position, whereas lys, leu, arg, trp, and phe were commonly found in the second position. Similarly, at the C-terminus, specific residues are favoured; for example, residues lys, arg, leu, and trp are present at most locations. In the case of GV ABPs at the N-terminus, the first position is favoured by gly, phe, lys, arg, ile and leu, whereas leu, trp, lys, ile, arg, and phe was often found in the second position. While the C-terminus is dominated by the residues lys, leu, arg, trp, phe, ile, and gly.

**Figure 5:**
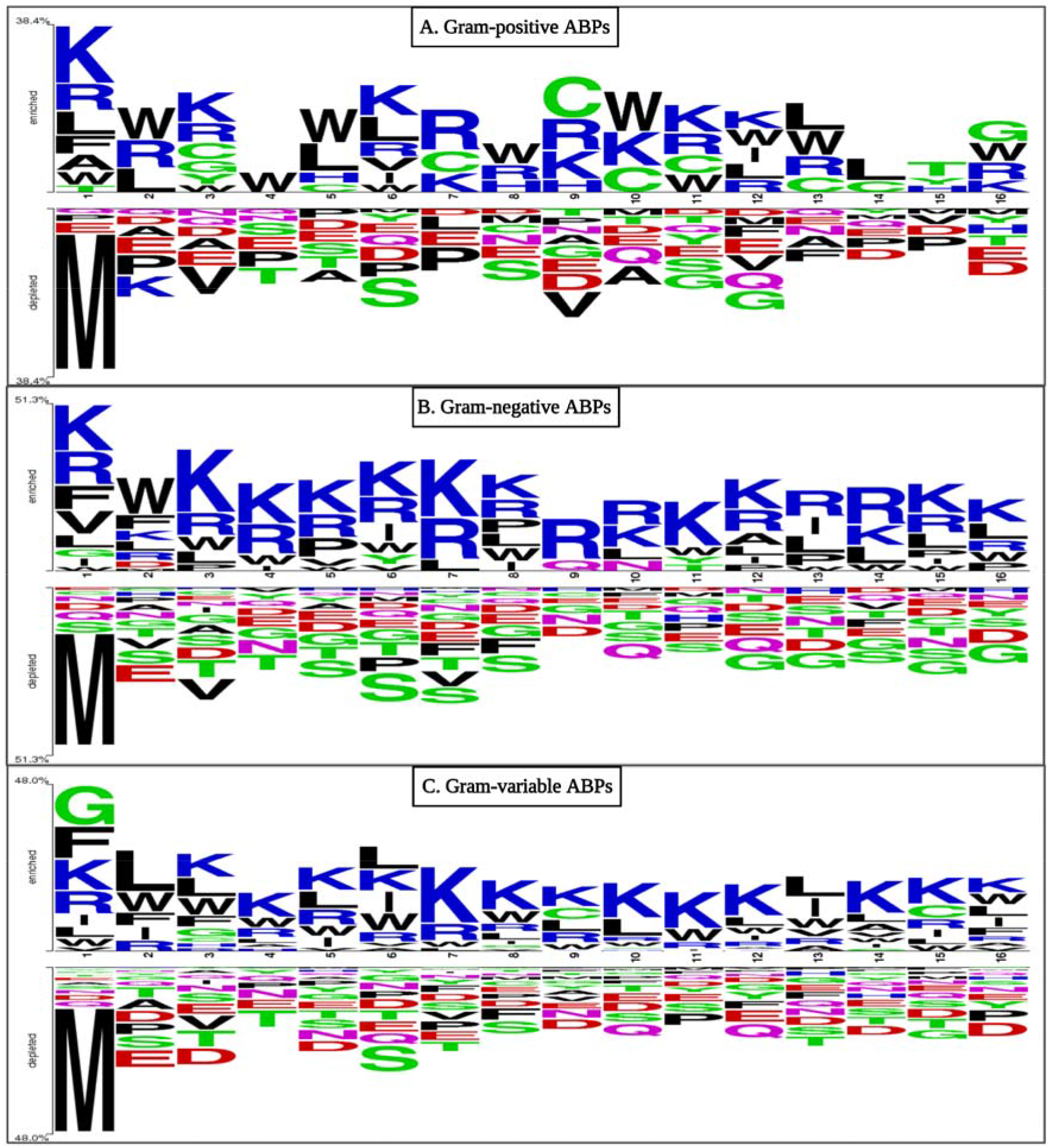
Two sample logo (TSL) representations of ABPs (A. Gram-positive, B. negative, and C. variable bacteria) showing preferred positions for amino acids. Here, the first eight residues belong to N-terminus, while the last eight residues belong to the C-terminus of the peptide.

In all three categories of ABPs, the N and C termini have a larger fraction of positively charged residues. The residue preference for non-ABPs reveals that met, glu, ala, ser, asp, asn, and gln predominate at the first position of the N-terminus. Overall, when the amino acid composition of ABP and non-ABP is compared, positively charged Lys and Arg are predominant in antibacterial peptides (see Figure 5). Similarly, Gly and Leu propensity are high in ABPs because they are hydrophobic in nature, which aids in bacterial lipid membrane integration.

### 3. Performance of BLAST-search

BLAST was used in the study to assign labels to three groups of antibacterial peptide sequences. We utilised BLAST to query the ABPs of the validation dataset against the three separate databases built from the three training groups of datasets in this approach. Various e-values were used to determine the optimal e-value at which the BLAST module performed the best. Hits were obtained from the total number of peptides in the validation set (372 for GP ABPs, 582 for GN ABPs and 3594 for GV ABPs). Results for e-value for three groups of ABPs from 1e-20 to 1e+3 for GP ABPs, GN ABPs and GV ABPs are shown below in Table 1. The BLAST module performs poorly at larger e-values, as BLAST allows random hits at higher e-values (see Table 1). As a result, while the number of hits grows with increasing e-values, performance suffers as it allows incorrect hits.

**Table 1:** Comparison of performance among BLAST modules at different e-values for GP, GN and GV ABPs.

| e-value | Hits |  |  | Correct hits |  |  | Incorrect hits |  |  | Percentage of correct hits |  |  |
| --- | --- | --- | --- | --- | --- | --- | --- | --- | --- | --- | --- | --- |
|  | GP | GN | GV | GP | GN | GV | GP | GN | GV | GP | GN | GV |
| **1.00E-20** | 39 | 30 | 470 | 39 | 28 | 450 | 0 | 2 | 20 | 100 | 93 | 96 |
| **1.00E-10** | 110 | 138 | 1188 | 104 | 133 | 1042 | 6 | 5 | 146 | 95 | 96 | 88 |
| **1.00E-06** | 150 | 234 | 1516 | 141 | 226 | 1301 | 9 | 8 | 215 | 94 | 97 | 86 |
| **0.0001** | 173 | 262 | 1730 | 162 | 253 | 1477 | 11 | 9 | 253 | 94 | 97 | 85 |
| **0.001** | 195 | 284 | 1925 | 181 | 275 | 1649 | 14 | 9 | 276 | 93 | 97 | 86 |
| **0.01** | 211 | 326 | 2114 | 196 | 315 | 1809 | 15 | 11 | 305 | 93 | 97 | 86 |
| **0.1** | 230 | 362 | 2326 | 211 | 349 | 1992 | 19 | 13 | 334 | 92 | 96 | 86 |
| **1** | 272 | 416 | 2608 | 242 | 395 | 2228 | 30 | 21 | 380 | 89 | 95 | 85 |
| **10** | 329 | 506 | 3222 | 268 | 462 | 2658 | 61 | 44 | 564 | 81 | 91 | 82 |
| **50** | 357 | 552 | 3488 | 286 | 489 | 2840 | 71 | 63 | 648 | 80 | 89 | 81 |
| **100** | 359 | 557 | 3537 | 289 | 492 | 2869 | 70 | 65 | 668 | 81 | 88 | 81 |
| **200** | 361 | 564 | 3560 | 290 | 498 | 2884 | 71 | 66 | 676 | 80 | 88 | 81 |
| **1000** | 366 | 571 | 3584 | 295 | 500 | 2902 | 71 | 71 | 682 | 81 | 88 | 81 |

#GP: gram-positive, GN: gram-negative, GV: gram-variable

### 4. Performance of Motif-based approach

Peptide motifs are known to affect the antibacterial properties and aid in the discovery of conserved domains; therefore, in this study, with this belief, we attempted to locate unique motifs in all three groups of ABPs. We utilized the MERCI program to extract the motifs present only in each of the three categories of antibacterial peptides in the training dataset. The training dataset was used to identify discriminatory patterns for each group, and the existence of these motifs was then utilized to give group labels to each peptide sequence in the validation dataset. The number of distinct motifs at three different frequencies in the training set and the proportion of accurate motifs discovered in the validation dataset for each group are shown in Table 2.

**Table 2:**
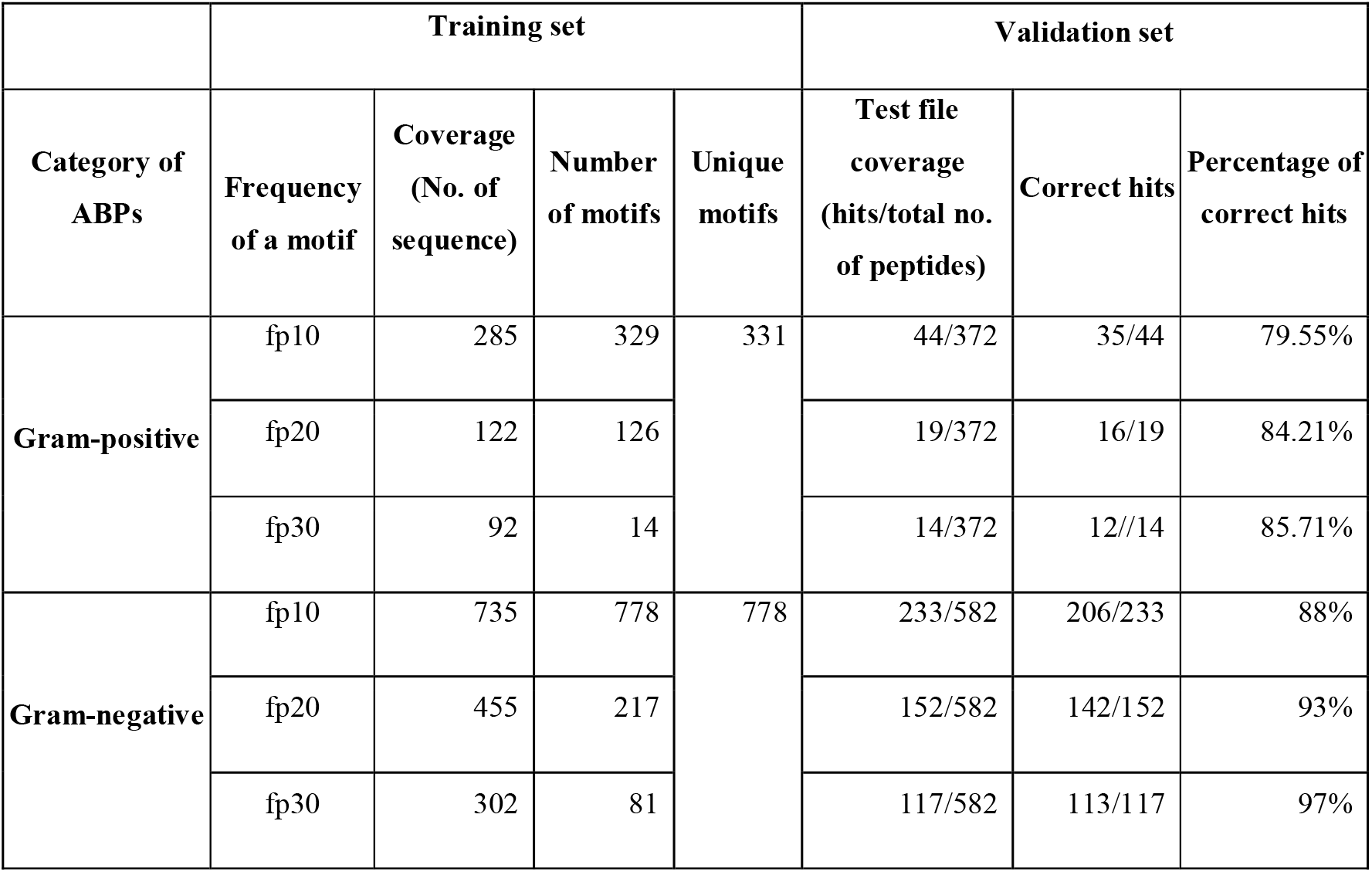
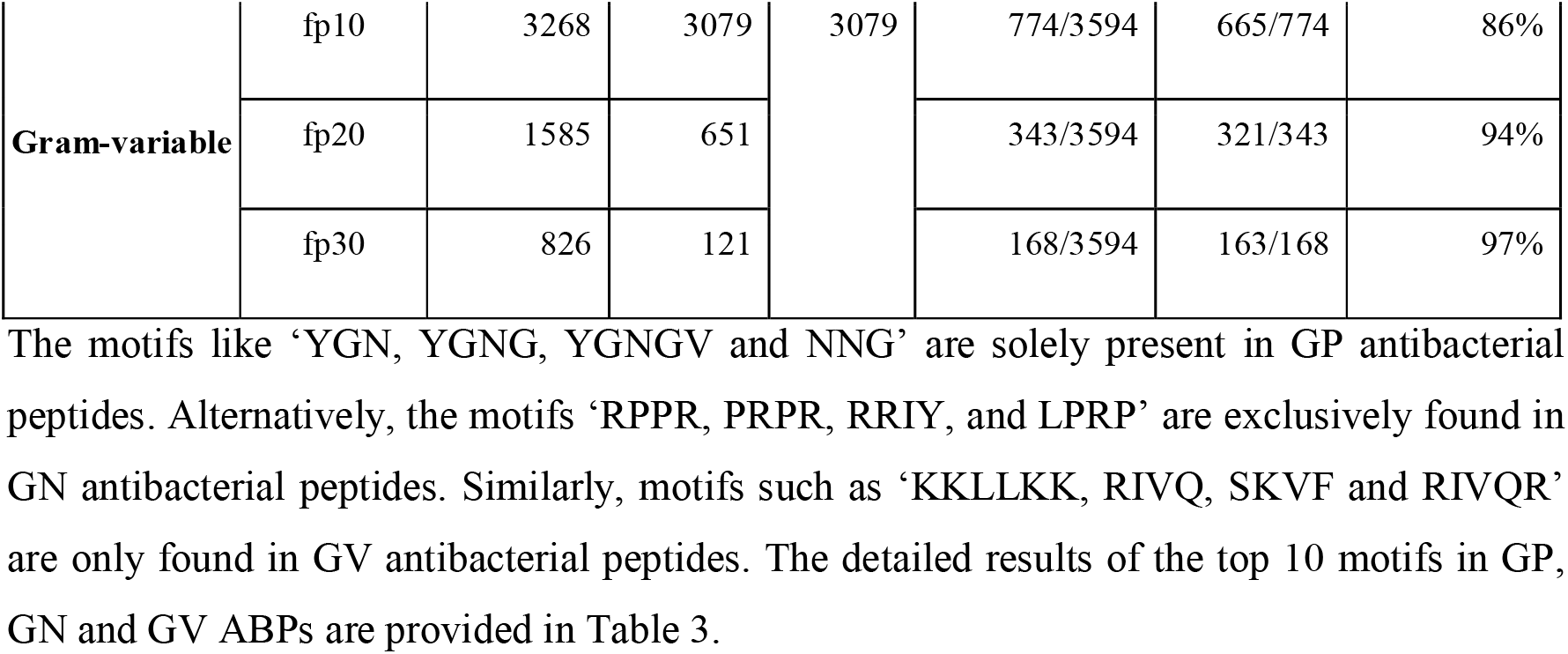
Number of unique motifs found in GP, GN and GV ABPs at three different frequencies in the training and validation dataset.

|  | Training set |  |  |  | Validation set |  |  |
| --- | --- | --- | --- | --- | --- | --- | --- |
| Category of ABPs | Frequency of a motif | Coverage (No. of sequence) | Number of motifs | Unique motifs | Test file coverage (hits/total no. of peptides) | Correct hits | Percentage of correct hits |
| Gram-positive | fp10 | 285 | 329 | 331 | 44/372 | 35/44 | 79.55% |
|  | fp20 | 122 | 126 |  | 19/372 | 16/19 | 84.21% |
|  | fp30 | 92 | 14 |  | 14/372 | 12//14 | 85.71% |
| Gram-negative | fp10 | 735 | 778 | 778 | 233/582 | 206/233 | 88% |
|  | fp20 | 455 | 217 |  | 152/582 | 142/152 | 93% |
|  | fp30 | 302 | 81 |  | 117/582 | 113/117 | 97% |
| **Gram-variable** | fp10 | 3268 | 3079 | 3079 | 774/3594 | 665/774 | 86% |
|  | fp20 | 1585 | 651 |  | 343/3594 | 321/343 | 94% |
|  | fp30 | 826 | 121 |  | 168/3594 | 163/168 | 97% |

The motifs like ‘YGN, YGNG, YGNGV and NNG’ are solely present in GP antibacterial peptides. Alternatively, the motifs ‘RPPR, PRPR, RRIY, and LPRP’ are exclusively found in GN antibacterial peptides. Similarly, motifs such as ‘KKLLKK, RIVQ, SKVF and RIVQR’ are only found in GV antibacterial peptides. The detailed results of the top 10 motifs in GP, GN and GV ABPs are provided in Table 3.

**Table 3:** Top 10 Motifs exclusively present in GP, GN and GV antibacterial peptide sequences.

| **S.no.** | **Gram-positive** |  | **Gram-negative** |  | **Gram-variable** |  |
| --- | --- | --- | --- | --- | --- | --- |
|  | **Motifs** | **Hits**<br>**(No. of**<br>**sequences)** | **Motifs** | **Hits**<br>**(No. of**<br>**sequences)** | **Motifs** | **Hits**<br>**(No. of**<br>**sequences)** |
| **1** | Y G N | 47 | R P P R | 78 | K K L L K K | 63 |
| **2** | Y G N G | 47 | P R P R | 74 | R I V Q | 54 |
| **3** | N N G | 44 | R R I Y | 71 | S K V F | 52 |
| **4** | Y G N G V | 38 | L P R P | 67 | R I V Q R | 51 |
| **5** | Y Y G N | 36 | I Y N | 67 | I V Q R I | 50 |
| **6** | Y Y G N G | 36 | I Y N R | 66 | K R I V Q | 49 |
| **7** | V D W | 34 | P R R I | 65 | V V I R | 48 |
| **8** | N G L P | 31 | P R R I Y | 65 | R W W R | 48 |
| **9** | P T G L | 30 | R I Y N | 65 | D F L R | 47 |
| **10** | R C R V | 30 | R R I Y N | 64 | R I V Q R I | 47 |

### 5. Machine Learning-based Models

We created three independent prediction models on training datasets of three groups of ABPs, utilizing different classifiers such as LR, RF, KNN, ET, SVC, DT, and GNB and analyzed the performance of these models on different feature sets.

#### 5.1. Performance of Composition-based Models

In this section, we computed the performance of 17 distinct compositional features. We discovered that ET-based classifiers outperformed all other classifiers by comparing the AUC-ROC curves (see Supplementary Figure S1). Using AAC-based features, we achieved maximum performance on the validation set with an AUC of 0.94 and MCC of 0.75 for GP ABPs. Similarly, employing PCP-based features on the validation set, we got comparable results (i.e., AUC = 0.94 and MCC = 0.72). Other composition-based features perform well across all three datasets. In the case of GN ABPs, we attain a maximum AUC of 0.98 and MCC of 0.86 using AAC-based features. On the validation set of GN ABPs, all other features likewise operate admirably. While using AAC-based features in the case of GV ABPs, we achieve an AUC of 0.95 and MCC of 0.74 on the validation set. Here, APAAC also performs better on the validation set, whereas SOC, BTC and SEP underperform. The complete results of the training and testing dataset on all features for GP, GN and GV ABPs are provided in Table 4, and the rest of the results are provided in the supplementary Table S1, S2 and S3.

**Table 4:** The performance of extra tree models developed using 17 types of composition-based features on training and validation datasets, evaluated on GP, GN and GV ABPs.

| Feature Type | Training set |  |  |  |  |  | Validation set |  |  |  |  |  |
| --- | --- | --- | --- | --- | --- | --- | --- | --- | --- | --- | --- | --- |
|  | GP |  | GN |  | GV |  | GP |  | GN |  | GV |  |
|  | AUC | MCC | AUC | MCC | AUC | MCC | AUC | MCC | AUC | MCC | AUC | MCC |
| **AAC** | 0.93 | 0.71 | 0.96 | 0.80 | 0.93 | 0.71 | **0.94** | 0.75 | **0.98** | 0.86 | **0.95** | 0.74 |
| **DPC** | 0.91 | 0.68 | 0.95 | 0.77 | 0.92 | 0.70 | 0.93 | 0.72 | 0.98 | 0.86 | 0.92 | 0.70 |
| **ATC** | 0.84 | 0.52 | 0.91 | 0.69 | 0.86 | 0.58 | 0.84 | 0.51 | 0.95 | 0.78 | 0.89 | 0.62 |
| **BTC** | 0.75 | 0.36 | 0.79 | 0.43 | 0.76 | 0.38 | 0.70 | 0.29 | 0.84 | 0.51 | 0.76 | 0.37 |
| **CTC** | 0.88 | 0.59 | 0.94 | 0.75 | 0.90 | 0.68 | 0.90 | 0.68 | 0.97 | 0.82 | 0.91 | 0.71 |
| **PCP** | 0.91 | 0.64 | 0.95 | 0.76 | 0.91 | 0.67 | 0.94 | 0.72 | 0.97 | 0.84 | 0.93 | 0.72 |
| **AAI** | 0.91 | 0.67 | 0.95 | 0.77 | 0.91 | 0.68 | 0.92 | 0.71 | 0.97 | 0.86 | 0.92 | 0.73 |
| **RRI** | 0.89 | 0.61 | 0.94 | 0.75 | 0.91 | 0.68 | 0.91 | 0.66 | 0.97 | 0.82 | 0.92 | 0.69 |
| **PRI** | 0.87 | 0.58 | 0.94 | 0.72 | 0.90 | 0.66 | 0.88 | 0.63 | 0.97 | 0.80 | 0.91 | 0.67 |
| **DDR** | 0.91 | 0.65 | 0.94 | 0.73 | 0.91 | 0.69 | 0.93 | 0.67 | 0.97 | 0.80 | 0.92 | 0.72 |
| **SEP** | 0.61 | 0.18 | 0.72 | 0.37 | 0.74 | 0.35 | 0.60 | 0.16 | 0.80 | 0.44 | 0.79 | 0.45 |
| **SER** | 0.92 | 0.67 | 0.96 | 0.78 | 0.92 | 0.71 | 0.92 | 0.69 | 0.97 | 0.86 | 0.93 | 0.72 |
| **SPC** | 0.90 | 0.64 | 0.94 | 0.76 | 0.90 | 0.67 | 0.92 | 0.71 | 0.96 | 0.81 | 0.92 | 0.71 |
| **PAAC** | 0.93 | 0.70 | 0.96 | 0.80 | 0.92 | 0.71 | 0.93 | 0.72 | 0.97 | 0.87 | 0.93 | 0.74 |
| **APAAC** | 0.93 | 0.70 | 0.96 | 0.79 | 0.92 | 0.71 | 0.93 | 0.72 | 0.98 | 0.87 | 0.95 | 0.75 |
| **QSO** | 0.92 | 0.70 | 0.96 | 0.79 | 0.92 | 0.71 | 0.93 | 0.73 | 0.97 | 0.84 | 0.93 | 0.73 |
| **SOC** | 0.66 | 0.20 | 0.76 | 0.39 | 0.61 | 0.15 | 0.64 | 0.27 | 0.79 | 0.43 | 0.64 | 0.20 |

#### 5.2. Performance of Binary Profile-based Features

Since apart from composition, the order of amino acids is also a significant aspect to consider; therefore, we built models based on binary profiles of peptides to incorporate information about frequency as well as the order of residues. For all three categories of ABPs, we generated binary features and developed models using the N, C, and NC terminals.

For different binary profile features in the GP ABPs dataset, the RF, SVC, and ET models outperform all other classifiers (See Table 5). On the training set, AAB performed best for the NC-terminal, with AUC and MCC of 0.91 and 0.64, respectively, and a similar trend was found in the validation set, with AUC and MCC of 0.93 and 0.66, respectively. (The complete result is provided in the supplementary Table S4, S5 and S6) Similarly, for the GN ABPs dataset, the ET and SVC models outperform all other classifiers. Here, all the binary features perform quite well and show the highest AUC of 0.98 on the validation set for the NC-terminal with an MCC of 0.86. For the GV ABPs dataset, the SVC and RF models perform best among all other classifiers for the binary profile features. A similar trend was found here also, as the NC-terminal performed best for binary features on the training set, with an AUC of 0.92, and also performed well on the validation set, with an AUC of 0.94. Therefore, the models developed using the binary profile features of the NC-terminal perform superior to individual N and C terminus, suggesting that both N and C-terminal are crucial in distinguishing gram-positive ABPs from non-ABPs.

**Table 5:** The performance of ML-models, developed using binary profiles-based features at different terminals on training and validation datasets, evaluated on GP, GN and GV ABPs.

| Feature type | Terminal | Training set |  |  |  |  |  | Validation set |  |  |  |  |  |
| --- | --- | --- | --- | --- | --- | --- | --- | --- | --- | --- | --- | --- | --- |
|  |  | AUC | MCC | AUC | MCC | AUC | MCC | AUC | MCC | AUC | MCC | AUC | MCC |
| Category of ABP(Model) |  | GP (RF) |  | GN (ET) |  | GV (SVC) |  | GP (RF) |  | GN (ET) |  | GV (SVC) |  |
| AAB | N | 0.90 | 0.62 | 0.94 | 0.74 | 0.92 | 0.70 | 0.92 | 0.67 | 0.97 | 0.85 | 0.93 | 0.73 |
|  | C | 0.87 | 0.59 | 0.92 | 0.68 | 0.89 | 0.64 | 0.86 | 0.55 | 0.95 | 0.79 | 0.89 | 0.63 |
|  | NC | 0.91 | 0.64 | 0.95 | 0.75 | 0.92 | 0.73 | **0.93** | 0.66 | **0.98** | 0.86 | **0.94** | 0.74 |
| Category of ABP(Model) |  | GP (SVC) |  | GN (SVC) |  | GV (RF) |  | GP (SVC) |  | GN (SVC) |  | GV (RF) |  |
| DPB | N | 0.88 | 0.59 | 0.93 | 0.72 | 0.48 | -0.10 | 0.89 | 0.63 | 0.97 | 0.80 | 0.86 | 0.52 |
|  | C | 0.85 | 0.54 | 0.91 | 0.67 | 0.88 | 0.59 | 0.85 | 0.53 | 0.95 | 0.77 | 0.86 | 0.56 |
|  | NC | 0.90 | 0.64 | 0.94 | 0.73 | 0.91 | 0.67 | 0.91 | 0.68 | 0.98 | 0.87 | 0.67 | 0.32 |
| Category of ABP(Model) |  | GP (ET) |  | GN (ET) |  | GV (SVC) |  | GP (ET) |  | GN (ET) |  | GV (SVC) |  |
| AIB | N | 0.89 | 0.59 | 0.94 | 0.74 | 0.92 | 0.71 | 0.91 | 0.65 | 0.97 | 0.85 | 0.93 | 0.69 |
|  | C | 0.88 | 0.56 | 0.92 | 0.69 | 0.89 | 0.65 | 0.88 | 0.57 | 0.96 | 0.76 | 0.89 | 0.64 |
|  | NC | 0.91 | 0.65 | 0.95 | 0.73 | 0.92 | 0.73 | 0.93 | 0.69 | 0.98 | 0.85 | 0.94 | 0.73 |
| Category of ABP(Model) |  | GP (RF) |  | GN (ET) |  | GV (SVC) |  | GP (RF) |  | GN (ET) |  | GV (SVC) |  |
| PCB | N | 0.90 | 0.64 | 0.94 | 0.72 | 0.91 | 0.70 | 0.92 | 0.63 | 0.97 | 0.84 | 0.93 | 0.73 |
|  | C | 0.88 | 0.61 | 0.91 | 0.67 | 0.88 | 0.62 | 0.86 | 0.57 | 0.94 | 0.74 | 0.88 | 0.63 |
|  | NC | 0.91 | 0.64 | 0.94 | 0.73 | 0.92 | 0.72 | 0.93 | 0.70 | 0.98 | 0.86 | 0.94 | 0.75 |

#### 5.3. Performance of word embedding features

In this feature extraction method, we wanted to assess the efficiency of different segmentation sizes and the n-gram mixture employed. We evaluated the overall performance of several classifiers from each ABPs group in both training and testing tests. The outcomes of the classifiers are shown in Table 6. Among the four segmentation sizes, the feature type corresponding to biological words of length = 1 and 2 contributed to the best overall average performance for all groups using an ET-based model. This demonstrates the significance of individual amino acid residues in an antibacterial peptide sequence. The complete results are provided in the supplementary Table S8.

**Table 6:** The performance of best ML models developed using FastText-based features on training and validation datasets, evaluated on GP, GN and GV ABPs.

| Feature<br>(n-gram<br>size) | Training set |  |  |  |  |  | Validation set |  |  |  |  |  |
| --- | --- | --- | --- | --- | --- | --- | --- | --- | --- | --- | --- | --- |
|  | GP |  | GN |  | GV |  | GP |  | GN |  | GV |  |
|  | AUC | MCC | AUC | MCC | AUC | MCC | AUC | MCC | AUC | MCC | AUC | MCC |
| **1g** | 0.91 | 0.67 | 0.95 | 0.77 | 0.89 | 0.64 | **0.92** | 0.68 | **0.97** | 0.84 | 0.92 | 0.71 |
| **2g** | 0.91 | 0.66 | 0.94 | 0.75 | 0.92 | 0.71 | 0.91 | 0.71 | 0.97 | 0.82 | **0.93** | 0.74 |
| **3g** | 0.87 | 0.57 | 0.92 | 0.70 | 0.91 | 0.68 | 0.84 | 0.56 | 0.96 | 0.82 | 0.92 | 0.68 |
| **2g & 3g**<br>**Combined** | 0.91 | 0.66 | 0.95 | 0.77 | 0.92 | 0.73 | 0.90 | 0.65 | 0.97 | 0.82 | 0.93 | 0.73 |

### 6. Performance of cross-prediction

We evaluated the performance of each binary features-based ML model in relation to the receiver operating characteristic (ROC) curve on both training and testing datasets, and the results are provided in a supplementary Figure S2. According to the ROC comparison, the RF-based model outperformed all other models in the case of GP ABPs, the ET-based model outperformed all other models in the case of GN ABPs, and the SVC-based model exceeded all other models in the case of GV ABPs for AAB binary feature at NC-terminal. As a result, in our further study, we applied RF-based, ET-based, and SVC-based models for GP, GN, and GV ABPs, respectively. Therefore, we have done a cross-prediction analysis of our best-performing models, and the performance of each of the final models on different validation datasets and the best performance is highlighted in bold, shown in Table 7. The models perform best with their own datasets and are significantly poor when making predictions with different groups of validation datasets. The AUC of the models falls with different sets of validation data, such as in the case of gram-positive ABPs, where the model trained with GP ABPs has an AUC of 0.93 on the GP validation dataset but drops to 0.89 and 0.91 on the GN and GV validation datasets. In the case of GN ABPs, models attain an AUC of 0.98 on their own validation dataset but drop to 0.85 and 0.93 on the GP and GV validation datasets, respectively. Similarly, the AUC of the GV ABPs model on its own validation dataset is 0.94, which drops to 0.88 and 0.93 for GP and GN validation datasets. This trend indicates the necessity for developing different models for each of the three groups of ABPs. We have also generated a confusion matrix of the best-performing model using the binary feature AAB (NC-terminal), which is given in the supplementary Figure S3. The visualization of the confusion matrix provides valuable insights into the model’s predictive accuracy and reveals areas for potential improvement.

**Table 7:** The analysis of the cross-prediction performance of RF, ET and SVC-based models for GP, GN and GV ABPs, respectively, using AAB binary feature of NC-terminal on different validation datasets.

| Validation dataset | Performance measures | Models |  |  |
| --- | --- | --- | --- | --- |
|  |  | Gram-positive | Gram-negative | Gram-variable |
| Gram-positive ABPs | Sn | 80.65 | 50.00 | 67.74 |
|  | Sp | 90.32 | 89.25 | 87.63 |
|  | Acc | 85.48 | 69.62 | 77.69 |
|  | AUC | **0.93** | 0.85 | 0.88 |
|  | AUPRC | 85.48 | 69.62 | 77.69 |
|  | MCC | 0.71 | 0.43 | 0.57 |
| Gram-negative ABPs | Sn | 65.98 | 92.44 | 91.07 |
|  | Sp | 89.35 | 94.85 | 87.29 |
|  | Acc | 77.66 | 93.64 | 89.18 |
|  | AUC | 0.89 | **0.98** | 0.93 |
|  | AUPRC | 0.85 | 0.98 | 0.92 |
|  | MCC | 0.57 | 0.87 | 0.78 |
| Gram-variable ABPs | Sn | 80.19 | 76.24 | 83.92 |
|  | Sp | 87.37 | 92.04 | 90.10 |
|  | Acc | 83.78 | 84.14 | 87.01 |
|  | **AUC** | 0.91 | 0.93 | **0.94** |
|  | **AUPRC** | 0.89 | 0.93 | 0.93 |
|  | **MCC** | 0.68 | 0.69 | 0.74 |

### 7. Performance of Hybrid Approaches

From the above results, we have observed that AAB-based features for the NC terminal outperformed for all three groups of ABPs prediction models. Therefore, to construct the final predictions, we coupled BLAST similarity and MOTIF scores with best-performing ML-model scores computed using AAB (NC-terminal) features.

#### 7.1. ML-based models with BLAST search

In this study, in order to improve the performance of the individual ML models, we developed three hybrid models using the BLAST + ML technique to categorise ABPs into three categories. We began by using a similarity search (BLAST) to predict positive and negative peptides. We calculated the performance of three hybrid models on validation datasets using the best feature and best model at different e-value cut-offs. Once the ML model has made the forecast, the presence of the hit is utilised to fine-tune the prediction.

In the case of GP ABPs, the performance of the hybrid approach with BLAST slightly increases than the individual ML approach with AUC and AUPRC of 0.94 and 0.93, respectively, at an e-value of 1e-20. However, there is no substantial increase in the performance of the hybrid model using this method in the other two scenarios for GN and GV ABPs, even at the lowest e-value of 1e-20 evaluated. This suggests that the hybrid approach with BLAST is inefficient in doing the classification job better than individual ML predictions. The detailed results of the hybrid model with this BLAST approach are provided in Supplementary Table S9, S10 and S11.

#### 7.2. ML-based models with motif approach

Similarly, here also, to improve the performance of the individual ML models, we constructed three hybrid models using the MOTIF-search (MERCI) + ML technique to classify ABPs into three categories. First, we utilised the MOTIF search to predict positive and negative peptides based on whether or not a motif was detected in validation data. The best model’s score is then added to the MOTIF score to calculate the performance of three hybrid models on validation datasets at varying frequencies. However, we observed the same pattern as with the BLAST+ML strategy, with no substantial improvement in the performance of the hybrid model. The detailed results of the hybrid model with this MOTIF approach are provided in Supplementary Table S12, S13 and S14.

Overall, combining ML-based models with blast and motif approaches does not outperform than individual ML-based models.

### 8. Comparison with Other Prediction Tools

Our best-performing models were compared to five different ML and DL-based ABP predictors, namely, AMPScanner vr.2 [18], AI4AMP [15], iAMPpred [20], AMPfun [16], and ABP-Finder [12]. Currently, only a few tools, such as ABP-Finder and AMPfun, examine the distinction of antibacterial peptide groups based on gram staining. Other techniques, such as AMPScanner vr2, iAMPpred, and AI4AMP, predict whether or not the peptide has broad antibacterial action. The validation set that we have created for each group of ABPs is utilised for external testing by performing the prediction job on their web servers to compare the performance with our model. In the case of the two-stage classifier ABP-Finder, the performance was evaluated based on the type of validation set employed as it considers classes of antibacterial peptides. Predictions of the validation set of GP ABPs are regarded correct only if designated as gram-positive; otherwise, predictions of gram-negative or both are considered incorrect. Similarly, the accurate estimation for GN ABPs should be gram-negative, whereas predictions given as gram-positive or both are incorrect. In the case of GV ABPs, showing predictions as both are regarded right, whereas either gram-positive or gram-negative prediction is considered incorrect. Because of this, we are obtaining the final output in binary form; therefore, AUC and AUPRC cannot be computed for ABB-finder as both rely on probability predictions. However, ABP-Finder predictions are extremely poor for gram-positive and gram-negative datasets and average for gram-variable ABPs despite it considering the classification of ABPs into different groups. Similarly, the discrepancy in sensitivity and specificity for all other tools is very large for all three sets of ABPs, as shown in Table 8, indicating that the methods can predict antibacterial peptides but cannot accurately categorise antibacterial peptides. When we ran the validation set on our model, we got AUCs of 0.93, 0.98, and 0.94 for GP ABPs, GN ABPs, and GV ABPs, which is greater than any of the abovementioned tools. Because of fewer false positives and the greatest AUC, AntiBP3 is able to attain a balanced sensitivity and specificity. The comprehensive comparison of prediction by web servers is provided in Table 8.

**Table 8:**
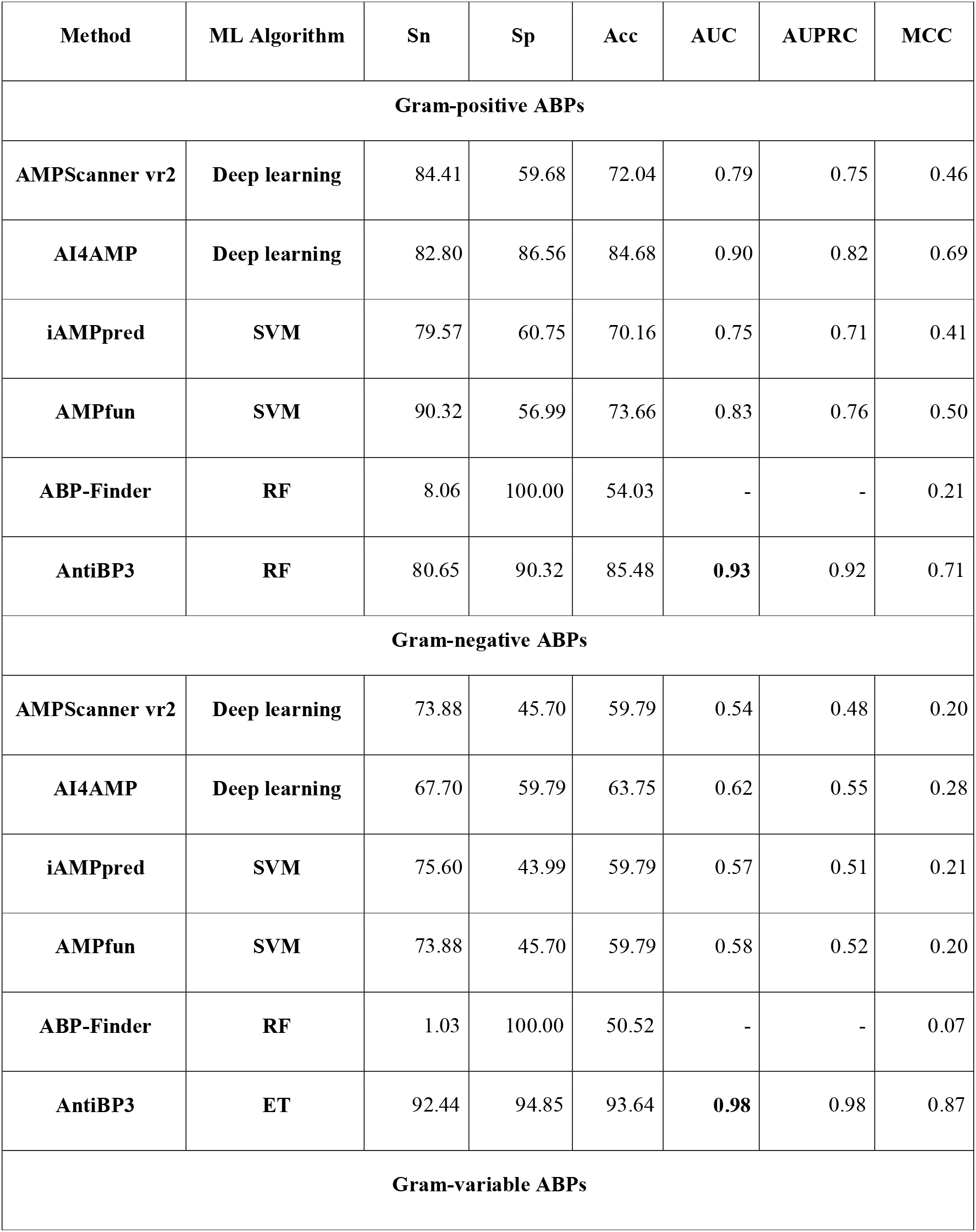
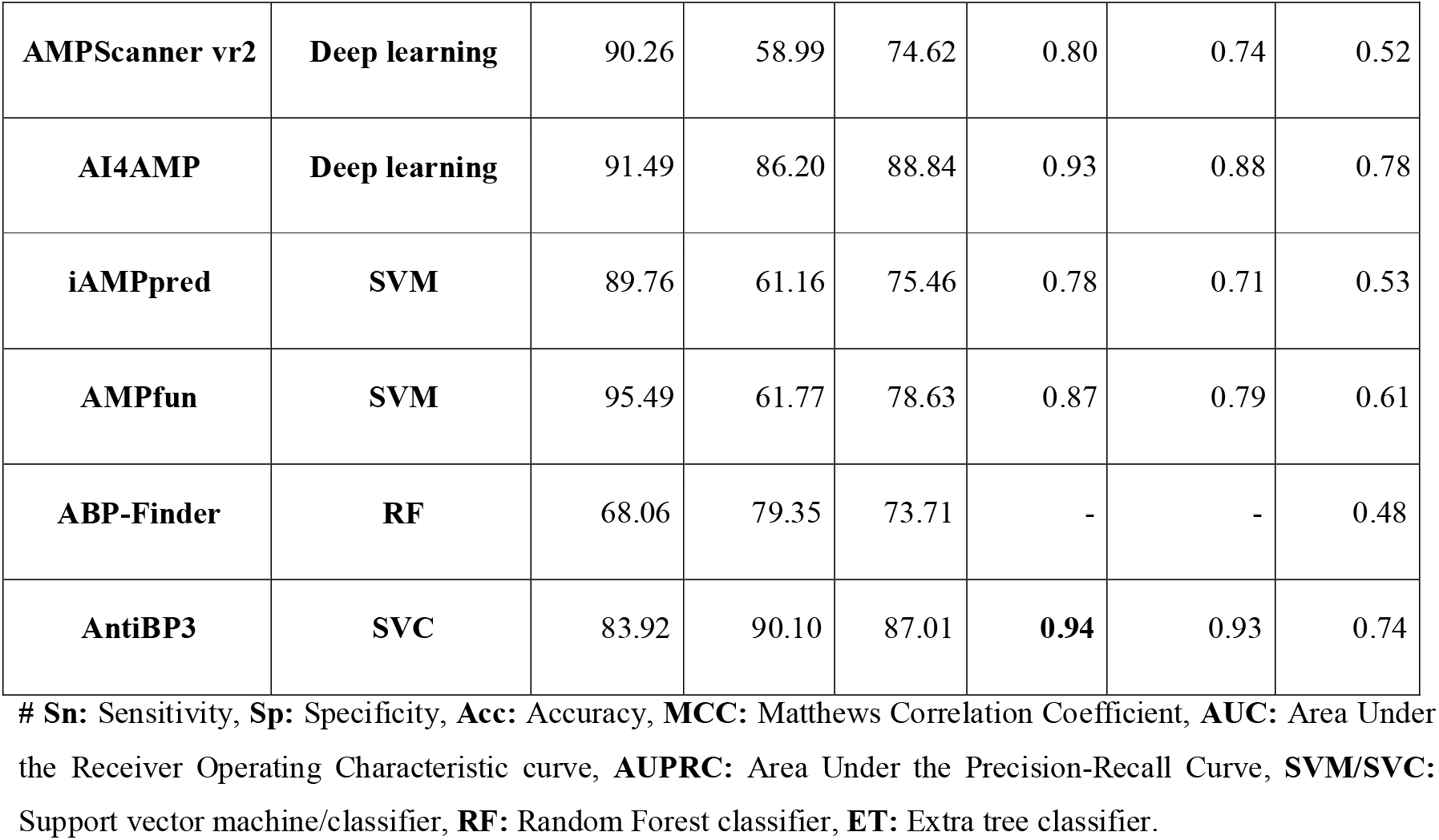
The comparison of existing prediction tools with AntiBP3 on the validation dataset to discriminate the three groups of ABPs, i.e., gram-positive, negative and variable.

### 9. Implementation of AntiBP3 server

Our emphasis on applying regulatory concepts to the development of ML-based predictors stems from our aim to provide a publicly available and well-maintained tool for reliably screening peptide libraries. To that purpose, we integrated our models in AntiBP3, a user-friendly online server (https://webs.iiitd.edu.in/raghava/antibp3/) that predicts Antibacterial peptides for different groups of bacteria based on sequence information for each entry. The web server includes the best prediction models for all three classes of ABPs. With a single submission operation, this programme can test hundreds of peptides effectively. The web server has been embraced with five main modules: (i) Predict, (ii) Design, (iii) BLAST scan, (iv) Motif scan, and (v) Package. Using a machine-learning method, the “Predict module” allows users to distinguish antibacterial peptides from non-antibacterial peptides for gram-positive, gram-negative, and gram-variable bacteria. The “Design module” enables the user to generate/design all potential mutants of the query sequence and predict whether or not the mutants exhibit antibacterial activity. The “BLAST scan module” employs a similarity search strategy to identify a hit from the input query sequence against the customised databases generated using known ABPs and non-ABPs for the three bacteria groups. The “Motif scan module” allows users to discover conserved sections in query sequences from MERCI-extracted motifs for the three antibacterial peptide groups. The ’AntiBP3’ server was created utilising several templates such as HTML, JAVA, and PHP, making it compatible with a variety of operating systems. The web server (https://webs.iiitd.edu.in/raghava/antibp3), python based standalone package (https://webs.iiitd.edu.in/raghava/antibp3/package.php) and GitHub (https://github.com/raghavagps/antibp3) are freely-accessible which will serve the scientific community to screen regions within a long amino acid sequence to identify promising antibacterial fragments for different bacterial species.

## Discussion

ABPs, belonging to the class of antimicrobials, play a crucial role in innate immunity similar to Defensins and exhibit potent activity against drug-resistant bacteria [53]. The field of in silico prediction of antibacterial peptides has witnessed the development of various methods, including AntiBP, ABP-Finder, Deep-ABPpred, StaBle-ABPpred, and AntiBP2. One of the major obstacles faced by researchers is the continuous updating of these models to incorporate the latest information. In 2010, we developed AntiBP2 for predicting ABPs, which was trained on a small dataset (999 ABPs and 999ABPs). Despite it being heavily used and cited by the scientific community, it is not updated in the last one decade. To address this issue, we presented AntiBP3, a method that is trained, tested, and evaluated on the largest possible data using the most up-to-date information. We opted to create three independent models to better accommodate the particular characteristics of different bacterial groups and improve the prediction accuracy. Rather than adopting a single generalised model like AntiBP2, each model was created to cater to a specific group of bacteria, allowing us to capture the distinct features and variations within each group. Universally, bacterial species are categorised as gram-positive or gram-negative based on the structural makeup of their cell walls as assessed by gram-staining, the gold standard staining method. A gram-positive bacterial cell’s surface is made up of a thick peptidoglycan layer that is rich in teichoic acid and lacks an outer membrane. The cell wall of a gram-negative bacterial cell consists of a thin peptidoglycan layer and an outer lipopolysaccharide membrane. In contrast, some bacterial species don’t belong to the two categories, such as *Mycobacterium* sp. or *Mycoplasma* sp., known as gram-indeterminate bacteria and some bacteria belonging to *Actinomyces* sp., *Bacillus sp. Clostridium* sp. shows an erratic response to the stain known as gram-variable bacteria [54]. However, considerable work has been done with respect to *Mycobacterium,* such as the creation of a database and prediction tools for AntiTB peptides, but not with other bacterial species [55, 56]. Thus, in this study, we created three prediction models to classify antibacterial peptides into three categories of ABP against gram-positive, gram-negative, and gram-variable bacteria.

We developed numerous prediction models using alignment-free ML techniques from sequence-based features, including compositional, binary profiles based on the standalone package of pfeature, and we also used fastText, a new emerging feature generation method in the field of antimicrobials. Using distinct features, the antibacterial activity of the peptides (ABPs or non-ABPs class) was predicted independently for each bacterial group. We investigated every possible feature in an attempt to predict the various categories of ABPs with high accuracy. We discovered that among all the compositional features-based models, the model created utilizing basic amino acid composition outperformed models developed using sophisticated features. In the case of binary-based features, the NC terminus, which considers both amino acid order and composition, performs better than N- or C-terminus-based features. This concordant with the trend followed by the AntiBP & AntiBP2 method; this suggests that both termini are important in imparting antibacterial properties to the peptide and enhancing its activity by interacting with the bacterial membrane with the C-terminal and the N-terminus later aids in modulating the critical intracellular process to kill the bacteria (see Table 5) [57, 58]. However, fastText features also perform comparably with compositional or binary-based features.

This research aims to answer an important biological question by developing specialized classification models that can reliably predict ABPs in datasets comprising gram-positive, gram-negative, and gram-variable bacteria. Consequently, we have done cross-prediction of the models on the validation set of different groups of ABPs, and the findings demonstrate a substantial decline in predicting labels for different datasets of ABPs when trained with another group of ABPs. In addition, we generated the motifs that are solely in GP ABPs, GN ABPs, and GV ABPs, which are listed in Table 3. Along with the ML approaches, commonly used alignment-based methods, BLAST and MOTIF-search, were used to annotate the peptide sequence. However, our hybrid models (BLAST + machine learning and MOTIF + machine learning) do not outperform individual ML-based models, which might be owing to the prevalence of false positives even in extreme cases such as at the lowest e-value of 1e-20, which reduces the AUC in hybrid techniques. As a result, only ML-based models are intended to predict which types of bacteria the peptides are active against. Finally, when compared to current ABP predictors, our method outperformed others with a balanced sensitivity and specificity and a high AUC for all three types of ABPs. This implies that our approach can more precisely categorise antibacterial peptides into their respective groups. These best-performing RF, ET, and SVC-based predictors for GP, GN, and GV ABPs, respectively, were implemented in the user-friendly and freely accessible web server AntiBP3, which was also used to identify new ABP hits from a large library of peptides for the scientific community, as well as a standalone package.

## Conclusion

Developing in-silico prediction tools for designing and synthesizing novel ABPs saves a significant amount of time and resources in screening peptide libraries. Separate prediction algorithms are required to address the relevance of AMPs’ particular activity, specifically antibacterial activity, source organism, and so on. In this study, we analyze different ML models that leverage distinct peptide properties for training. However, we demonstrated that all feature generation methods performed similarly. This highlighted that the crucial information for distinguishing antibacterial peptides can be captured solely from amino-acid sequence data without the need for sophisticated feature extraction methods. This will simplify and improve the interpretability of the modelling procedures. This study will aid in developing better and more effective antibacterial peptides against resistant strains of all bacterial classes in the future.

## Authors’ contributions

NB, AD and GPSR collected and processed the datasets. NB implemented the algorithms and developed the prediction models. NB and GPSR analyzed the results. SC created the back-end of the web server, and SC and NB created the from-end user interface. NB and GPSR performed the writing, reviewing and draft preparation of the manuscript. GPSR conceived and coordinated the project and gave overall supervision to the project. All authors have read and approved the manuscript.

## Acknowledgements

Authors are thankful to the Council of Scientific & Industrial Research (CSIR) and Department of Science and Technology (DST-INSPIRE) for fellowships and financial support and the Department of Computational Biology, IIITD New Delhi for infrastructure and facilities.

