## Supplementary file for "AntiBP3: A hybrid method for predicting antibacterial peptides against gram-positive/negative/variable bacteria"

Prof. Gajendra P. S. Raghava

Head and Professor

Department of Computational Biology

Indraprastha Institute of Information Technology, Delhi

Okhla Industrial Estate, Phase III (Near Govind Puri Metro Station)

New Delhi, India – 110020 Office: A-302 (R&D Block)

Website: <http://webs.iiitd.edu.in/raghava/>

**Supplementary Data**

**Supplementary Figure S1: AUC-ROC comparison of different ML models on Compositional based AAC features for GP, GN and GV ABPs.**

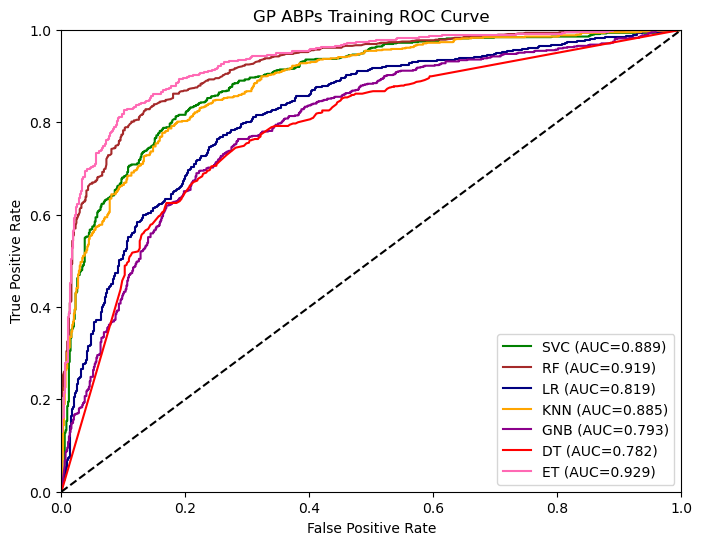

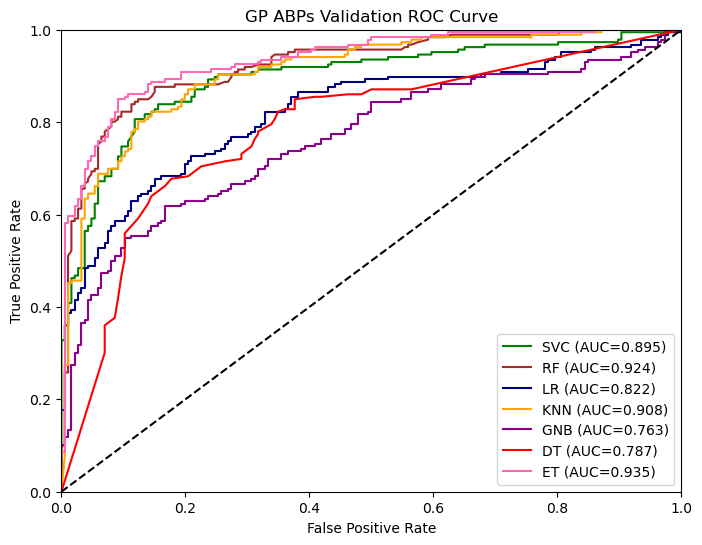

**Figure S1.1: The performance of ML models on training and testing data of GP ABPs on AAC feature**

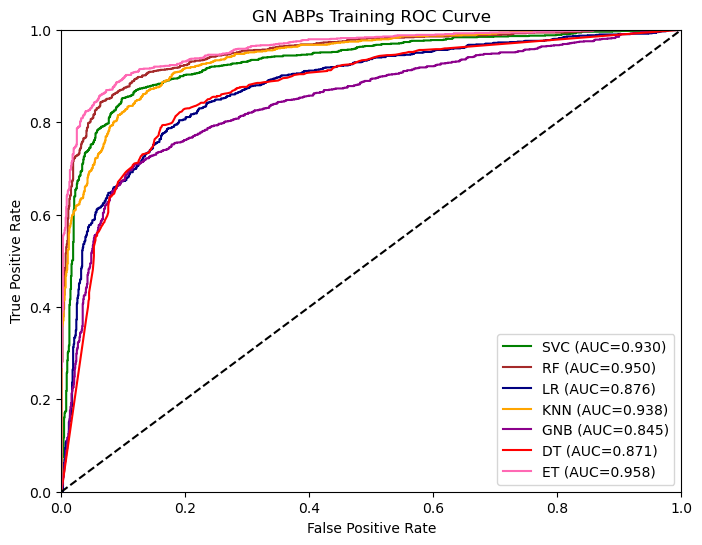

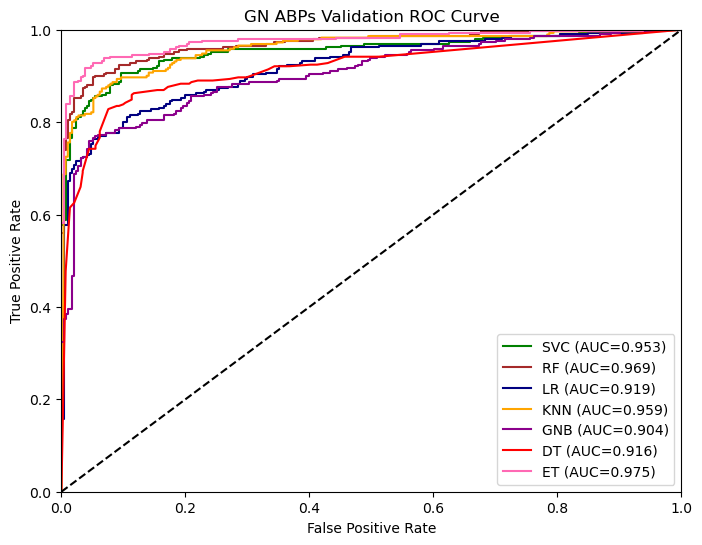

**Figure S1.2: The performance of ML models on training and testing data of GN ABPs on AAC feature**

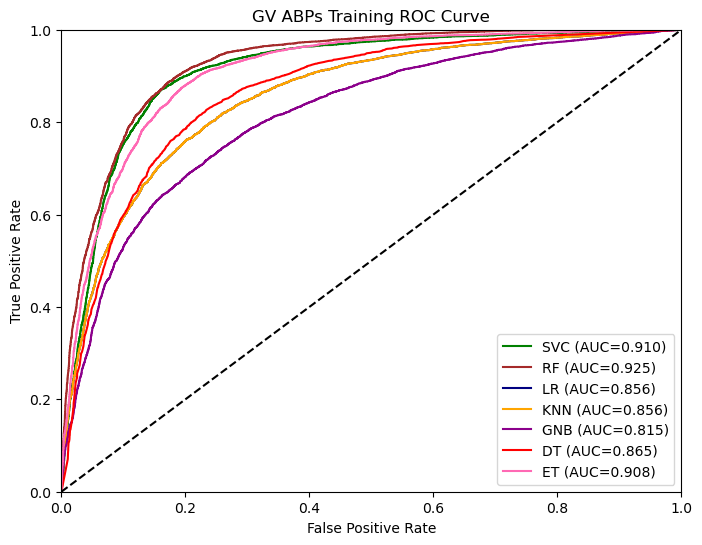

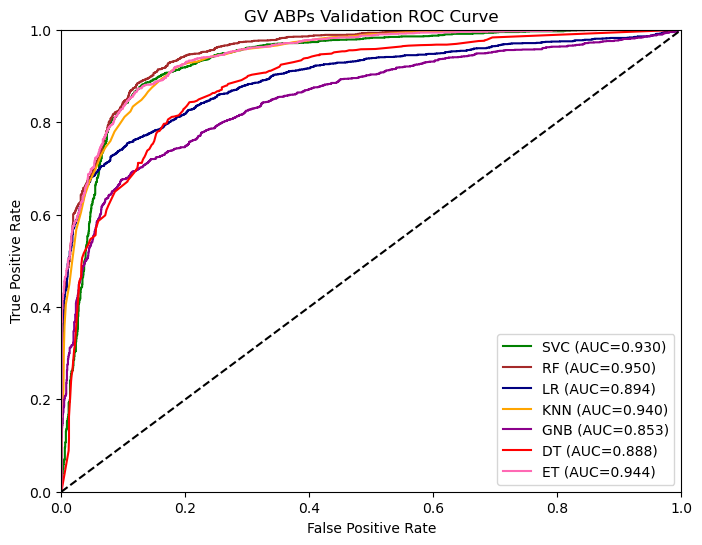

**Figure S1.3: The performance of ML models on training and testing data of GV ABPs on AAC feature**

**Supplementary Figure S2: The AUC-ROC comparison of different ML models on Binary-profile AAB features for NC-terminal for GP, GN and GV ABPs.**

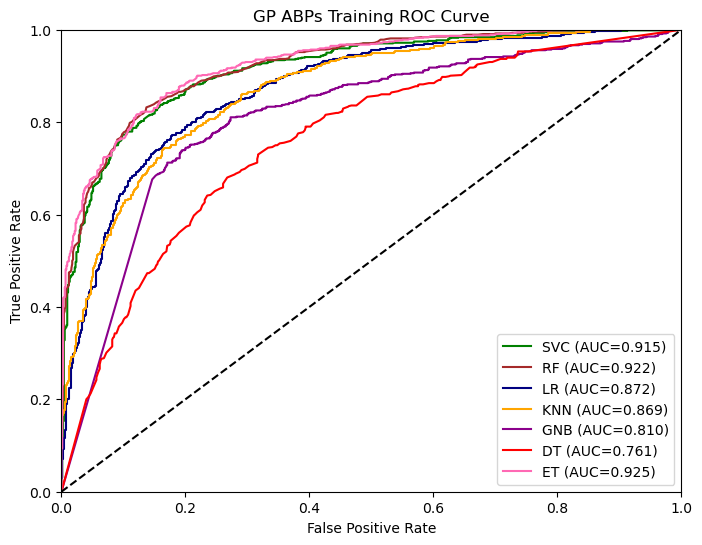

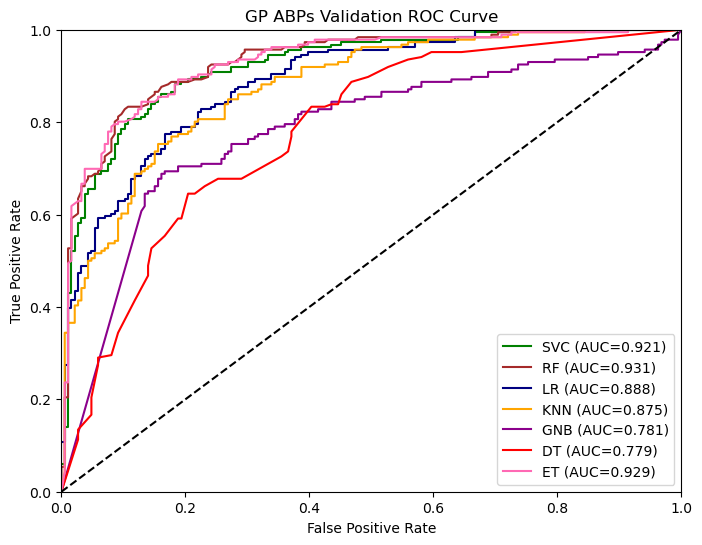

**Figure S2.1: The performance of ML models on training and testing data of GP ABPs on AAB (NC-terminal)**

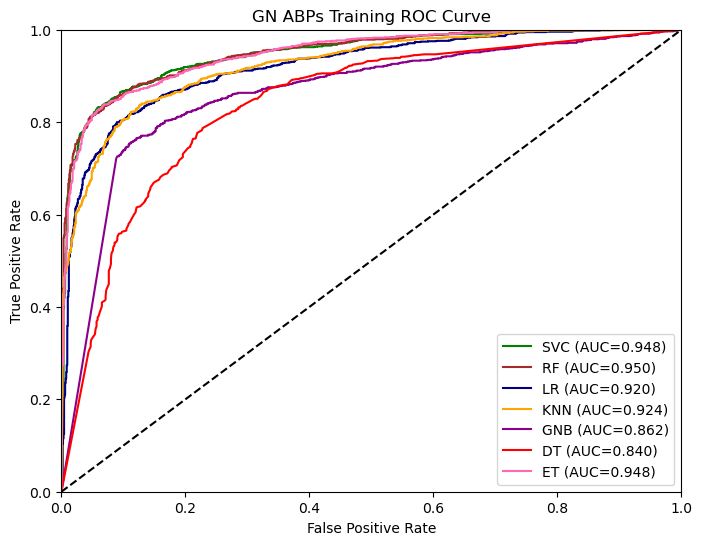

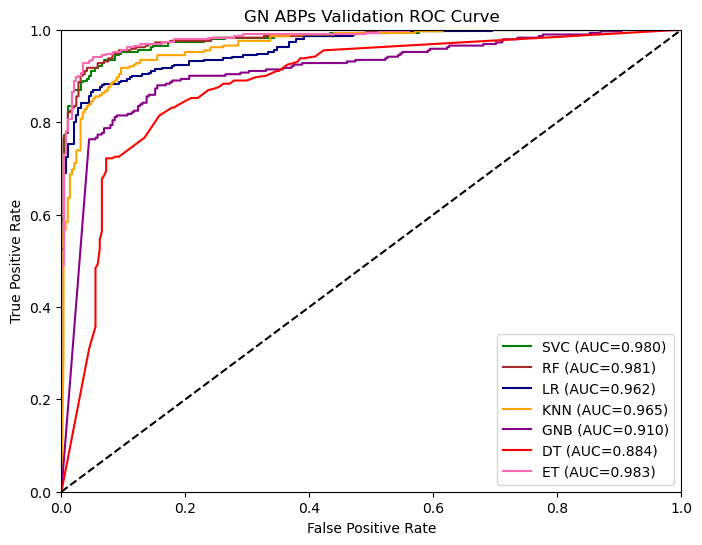

**Figure S2.2: Performance of ML models on training and testing data of GN ABPs on AAB (NC-terminal)**

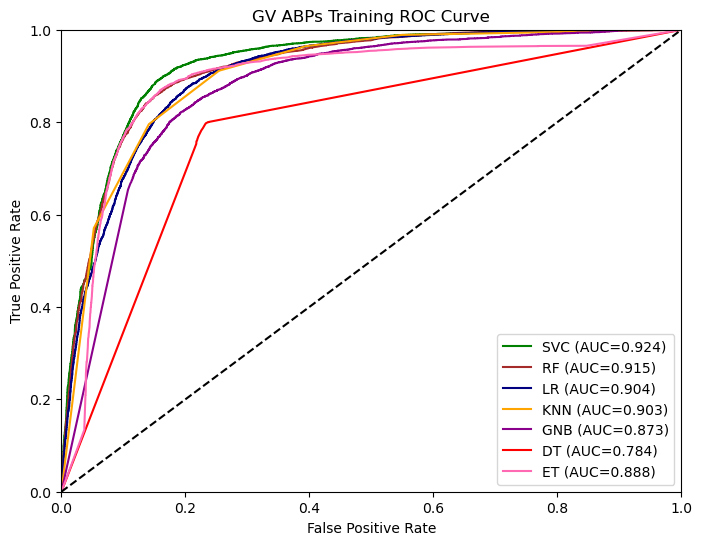

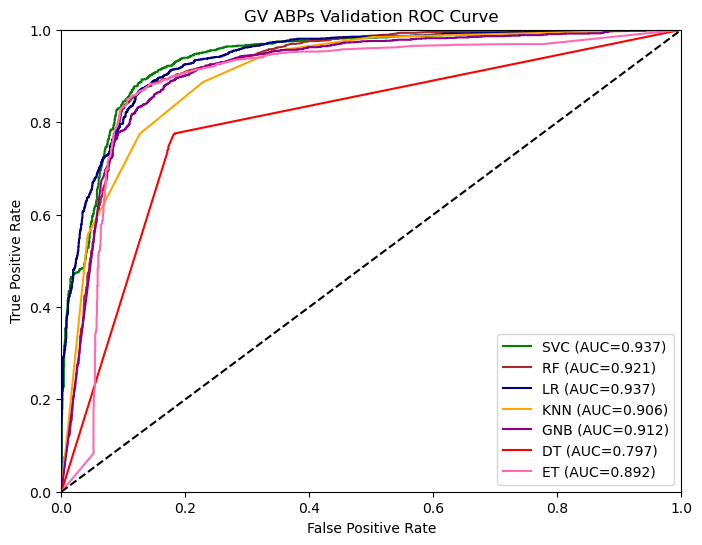

**Figure S2.3: Performance of ML models on training and testing data of GV ABPs on AAB (NC-terminal)**

**Supplementary Figure S3: Confusion matrix of best ML-model(AAB-based binary feature for NC-terminal) on the validation set of GP, GN and GV ABPs independent**

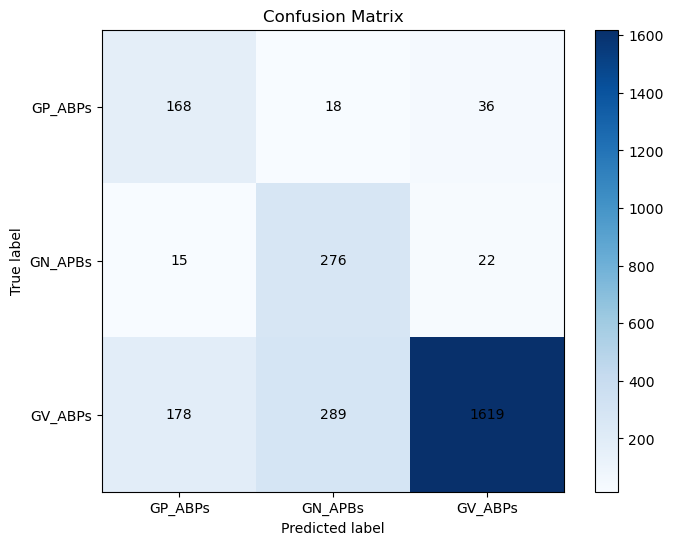

**Figure 7: Confusion matrix of all three groups of ABPs with the best ML model**

**Supplementary Tabel S1: Performance of Extra tree classifier on training and validation dataset developed using 17 types of composition-based features for gram-positive ABPs**

| **Feature Type** | **Training set** | | | | | | **Validation set** | | | | | |
| --- | --- | --- | --- | --- | --- | --- | --- | --- | --- | --- | --- | --- |
|  | **Sn** | **Sp** | **Acc** | **AUC** | **AUPRC** | **MCC** | **Sn** | **Sp** | **Acc** | **AUC** | **AUPRC** | **MCC** |
| **AAC** | 85.08 | 85.75 | 85.42 | 0.93 | 0.93 | 0.71 | 83.87 | 90.86 | 87.37 | 0.94 | 0.94 | 0.75 |
| **DPC** | 83.87 | 84.01 | 83.94 | 0.91 | 0.92 | 0.68 | 83.33 | 88.71 | 86.02 | 0.93 | 0.92 | 0.72 |
| **ATC** | 75.94 | 75.54 | 75.74 | 0.84 | 0.84 | 0.52 | 67.74 | 82.26 | 75.00 | 0.84 | 0.83 | 0.51 |
| **BTC** | 67.88 | 67.74 | 67.81 | 0.75 | 0.74 | 0.36 | 65.05 | 63.44 | 64.25 | 0.70 | 0.70 | 0.29 |
| **CTC** | 79.97 | 79.03 | 79.50 | 0.88 | 0.88 | 0.59 | 81.72 | 86.56 | 84.14 | 0.90 | 0.90 | 0.68 |
| **PCP** | 81.72 | 81.99 | 81.86 | 0.91 | 0.91 | 0.64 | 82.80 | 89.25 | 86.02 | 0.94 | 0.93 | 0.72 |
| **AAI** | 83.47 | 83.74 | 83.60 | 0.91 | 0.91 | 0.67 | 81.18 | 89.25 | 85.22 | 0.92 | 0.92 | 0.71 |
| **RRI** | 80.91 | 79.84 | 80.38 | 0.89 | 0.90 | 0.61 | 80.11 | 85.48 | 82.80 | 0.91 | 0.91 | 0.66 |
| **PRI** | 78.76 | 78.90 | 78.83 | 0.87 | 0.88 | 0.58 | 0.58 | 76.34 | 81.45 | 0.88 | 0.87 | 0.63 |
| **DDR** | 82.39 | 82.12 | 82.26 | 0.91 | 0.91 | 0.65 | 79.57 | 87.63 | 83.60 | 0.93 | 0.91 | 0.67 |
| **SEP** | 61.96 | 55.78 | 58.87 | 0.61 | 0.58 | 0.18 | 52.15 | 63.98 | 58.07 | 0.60 | 0.59 | 0.16 |
| **SER** | 83.87 | 83.33 | 83.60 | 0.92 | 0.92 | 0.67 | 78.50 | 90.32 | 84.41 | 0.92 | 0.92 | 0.69 |
| **SPC** | 82.26 | 81.45 | 81.86 | 0.90 | 0.90 | 0.64 | 81.18 | 89.25 | 85.22 | 0.92 | 0.91 | 0.71 |
| **PAAC** | 84.54 | 85.22 | 84.88 | 0.93 | 0.93 | 0.70 | 81.72 | 90.32 | 86.02 | 0.93 | 0.93 | 0.72 |
| **APAAC** | 85.48 | 84.68 | 85.08 | 0.93 | 0.93 | 0.70 | 83.87 | 88.17 | 86.02 | 0.93 | 0.92 | 0.72 |
| **QSO** | 84.68 | 85.08 | 84.88 | 0.92 | 0.92 | 0.70 | 82.80 | 90.32 | 86.56 | 0.93 | 0.92 | 0.73 |
| **SOC** | 59.95 | 59.95 | 59.95 | 0.66 | 0.67 | 0.20 | 58.60 | 67.74 | 63.17 | 0.64 | 0.63 | 0.27 |

**# Sn:** Sensitivity, **Sp:** Specificity, **Acc:** Accuracy, **MCC:** Matthews Correlation Coefficient, **AUC:** Area Under the Receiver Operating Characteristic curve, **AUPRC:**Area Under the Precision-Recall Curve, **AAC:** Amino acid composition, **APAAC:** Amphiphilic pseudo amino acid composition, **DDR:** Distance distribution of residue, **DPC:** Di-peptide composition, **QSO:** Quasi-sequence order, **PCP:** Physico-chemical properties composition, **PAAC:** Pseudo amino acid composition, **RRI:** Residue repeat Information, **SPC:** Shannon entropy of physicochemical properties, **ATC:** Atomic composition , **BTC:** Bond type composition, **CTC:** Conjoint triad descriptors, **AAI:** Amino Acid index, **PRI:** Property repeats index, **SEP:** Shannon entropy of a protein, **SER:** Shannon entropy of a residue, **SOC:** Sequence order coupling number

**Supplementary Tabel S2: Performance of Extra-tree classifier on training and validation dataset developed using 17 types of composition-based features for gram-negative ABPs**

| **Feature Type** | **Training set** | | | | | | **Validation set** | | | | | |
| --- | --- | --- | --- | --- | --- | --- | --- | --- | --- | --- | --- | --- |
|  | **Sn** | **Sp** | **Acc** | **AUC** | **AUPRC** | **MCC** | **Sn** | **Sp** | **Acc** | **AUC** | **AUPRC** | **MCC** |
| **AAC** | 90.03 | 89.61 | 89.82 | 0.96 | 0.96 | 0.80 | 93.13 | 93.13 | 93.13 | 0.98 | 0.98 | 0.86 |
| **DPC** | 88.23 | 88.32 | 88.27 | 0.95 | 0.96 | 0.77 | 94.50 | 91.41 | 92.96 | 0.98 | 0.98 | 0.86 |
| **ATC** | 84.11 | 84.45 | 84.28 | 0.91 | 0.92 | 0.69 | 88.66 | 89.69 | 89.18 | 0.95 | 0.96 | 0.78 |
| **BTC** | 71.39 | 71.13 | 71.26 | 0.79 | 0.79 | 0.43 | 76.63 | 74.57 | 75.60 | 0.84 | 0.86 | 0.51 |
| **CTC** | 87.46 | 87.97 | 87.72 | 0.94 | 0.94 | 0.75 | 92.10 | 90.03 | 91.07 | 0.97 | 0.98 | 0.82 |
| **PCP** | 88.32 | 87.89 | 88.10 | 0.95 | 0.95 | 0.76 | 91.07 | 92.78 | 91.92 | 0.97 | 0.98 | 0.84 |
| **AAI** | 88.49 | 87.97 | 88.23 | 0.95 | 0.95 | 0.77 | 92.44 | 93.13 | 92.78 | 0.97 | 0.98 | 0.86 |
| **RRI** | 87.54 | 87.20 | 87.37 | 0.94 | 0.95 | 0.75 | 89.00 | 93.13 | 91.07 | 0.97 | 0.97 | 0.82 |
| **PRI** | 86.34 | 85.83 | 86.08 | 0.94 | 0.94 | 0.72 | 90.03 | 89.69 | 89.86 | 0.97 | 0.97 | 0.80 |
| **DDR** | 86.68 | 86.68 | 86.68 | 0.94 | 0.95 | 0.73 | 88.32 | 91.41 | 89.86 | 0.97 | 0.97 | 0.80 |
| **SEP** | 70.02 | 66.67 | 68.34 | 0.72 | 0.68 | 0.37 | 69.76 | 75.60 | 73.65 | 0.80 | 0.63 | 0.44 |
| **SER** | 89.18 | 88.92 | 89.05 | 0.96 | 0.96 | 0.78 | 91.07 | 94.85 | 92.96 | 0.97 | 0.98 | 0.86 |
| **SPC** | 87.63 | 87.89 | 87.76 | 0.94 | 0.94 | 0.76 | 90.72 | 90.03 | 90.38 | 0.96 | 0.97 | 0.81 |
| **PAAC** | 90.12 | 89.78 | 89.95 | 0.96 | 0.96 | 0.80 | 92.78 | 94.16 | 93.47 | 0.97 | 0.98 | 0.87 |
| **APAAC** | 89.43 | 89.61 | 89.52 | 0.96 | 0.96 | 0.79 | 92.10 | 94.50 | 93.30 | 0.98 | 0.98 | 0.87 |
| **QSO** | 89.18 | 89.43 | 89.30 | 0.96 | 0.96 | 0.79 | 90.72 | 93.47 | 92.10 | 0.97 | 0.98 | 0.84 |
| **SOC** | 69.42 | 69.33 | 69.37 | 0.76 | 0.76 | 0.39 | 72.17 | 70.45 | 71.31 | 0.79 | 0.80 | 0.43 |

**# Sn:** Sensitivity, **Sp:** Specificity, **Acc:** Accuracy, **MCC:** Matthews Correlation Coefficient, **AUC:** Area Under the Receiver Operating Characteristic curve, **AUPRC:**Area Under the Precision-Recall Curve, **AAC:** Amino acid composition, **APAAC:** Amphiphilic pseudo amino acid composition, **DDR:** Distance distribution of residue, **DPC:** Di-peptide composition, **QSO:** Quasi-sequence order, **PCP:** Physico-chemical properties composition, **PAAC:** Pseudo amino acid composition, **RRI:** Residue repeat Information, **SPC:** Shannon entropy of physicochemical properties, **ATC:** Atomic composition , **BTC:** Bond type composition, **CTC:** Conjoint triad descriptors, **AAI:** Amino Acid index, **PRI:** Property repeats index, **SEP:** Shannon entropy of a protein, **SER:** Shannon entropy of a residue, **SOC:** Sequence order coupling number

**Supplementary Tabel S3: Performance of Extra-tree classifier on training and validation dataset developed using 17 types of composition-based features for gram-variable ABPs**

| **Feature Type** | **Training set** | | | | | | **Validation set** | | | | | |
| --- | --- | --- | --- | --- | --- | --- | --- | --- | --- | --- | --- | --- |
|  | **Sn** | **Sp** | **Acc** | **AUC** | **AUPRC** | **MCC** | **Sn** | **Sp** | **Acc** | **AUC** | **AUPRC** | **MCC** |
| **AAC** | 85.10 | 85.59 | 85.34 | 0.93 | 0.91 | 0.71 | 84.70 | 89.65 | 87.17 | 0.95 | 0.95 | 0.74 |
| **DPC** | 85.21 | 84.93 | 85.07 | 0.92 | 0.90 | 0.70 | 82.97 | 87.31 | 85.14 | 0.92 | 0.90 | 0.70 |
| **ATC** | 79.01 | 79.01 | 79.01 | 0.86 | 0.85 | 0.58 | 80.69 | 81.41 | 81.05 | 0.89 | 0.89 | 0.62 |
| **BTC** | 68.43 | 69.09 | 68.76 | 0.76 | 0.74 | 0.38 | 66.67 | 70.62 | 68.64 | 0.76 | 0.73 | 0.37 |
| **CTC** | 83.90 | 83.79 | 83.85 | 0.90 | 0.89 | 0.68 | 83.19 | 87.31 | 85.25 | 0.91 | 0.90 | 0.71 |
| **PCP** | 83.56 | 83.51 | 83.54 | 0.91 | 0.90 | 0.67 | 84.31 | 87.37 | 85.84 | 0.93 | 0.91 | 0.72 |
| **AAI** | 84.13 | 83.71 | 83.92 | 0.91 | 0.89 | 0.68 | 85.14 | 87.31 | 86.23 | 0.92 | 0.90 | 0.73 |
| **RRI** | 83.99 | 84.24 | 84.11 | 0.91 | 0.90 | 0.68 | 82.58 | 86.53 | 84.56 | 0.92 | 0.90 | 0.69 |
| **PRI** | 83.14 | 83.22 | 83.18 | 0.90 | 0.89 | 0.66 | 81.91 | 85.36 | 83.64 | 0.91 | 0.89 | 0.67 |
| **DDR** | 84.31 | 84.35 | 84.33 | 0.91 | 0.90 | 0.69 | 84.14 | 87.87 | 86.00 | 0.92 | 0.90 | 0.72 |
| **SEP** | 67.64 | 67.14 | 67.39 | 0.74 | 0.73 | 0.35 | 72.51 | 72.68 | 72.59 | 0.79 | 0.78 | 0.45 |
| **SER** | 85.23 | 85.43 | 85.33 | 0.92 | 0.91 | 0.71 | 84.14 | 88.20 | 86.17 | 0.93 | 0.91 | 0.72 |
| **SPC** | 83.50 | 83.03 | 83.26 | 0.90 | 0.89 | 0.67 | 83.53 | 87.31 | 85.42 | 0.92 | 0.91 | 0.71 |
| **PAAC** | 85.39 | 85.06 | 85.23 | 0.92 | 0.91 | 0.71 | 85.25 | 88.26 | 86.76 | 0.93 | 0.91 | 0.74 |
| **APAAC** | 85.62 | 85.38 | 85.50 | 0.92 | 0.91 | 0.71 | 85.42 | 89.65 | 87.54 | 0.95 | 0.95 | 0.75 |
| **QSO** | 85.31 | 85.45 | 85.38 | 0.92 | 0.90 | 0.71 | 84.25 | 88.43 | 86.34 | 0.93 | 0.91 | 0.73 |
| **SOC** | 57.35 | 57.22 | 57.28 | 0.61 | 0.62 | 0.15 | 59.88 | 59.66 | 59.77 | 0.64 | 0.65 | 0.20 |

**# Sn:** Sensitivity, **Sp:** Specificity, **Acc:** Accuracy, **MCC:** Matthews Correlation Coefficient, **AUC:** Area Under the Receiver Operating Characteristic curve, **AUPRC:**Area Under the Precision-Recall Curve, **AAC:** Amino acid composition, **APAAC:** Amphiphilic pseudo amino acid composition, **DDR:** Distance distribution of residue, **DPC:** Di-peptide composition, **QSO:** Quasi-sequence order, **PCP:** Physico-chemical properties composition, **PAAC:** Pseudo amino acid composition, **RRI:** Residue repeat Information, **SPC:** Shannon entropy of physicochemical properties, **ATC:** Atomic composition , **BTC:** Bond type composition, **CTC:** Conjoint triad descriptors, **AAI:** Amino Acid index, **PRI:** Property repeats index, **SEP:** Shannon entropy of a protein, **SER:** Shannon entropy of a residue, **SOC:** Sequence order coupling number

**Supplementary Tabel S4: Performance of ML-models on training and validation dataset developed using four types of Binary profile-based features of N, C, NC-terminal of gram-positive ABPs**

| **Feature type** | **Terminal** | **Training set** | | | | | | **Validation set** | | | | | |
| --- | --- | --- | --- | --- | --- | --- | --- | --- | --- | --- | --- | --- | --- |
|  |  | **Sn** | **Sp** | **Acc** | **AUC** | **AUPRC** | **MCC** | **Sn** | **Sp** | **Acc** | **AUC** | **AUPRC** | **MCC** |
| **AAB**  **(RF)** | **N** | 81.32 | 80.91 | 81.12 | 0.90 | 0.91 | 0.62 | 77.96 | 88.17 | 83.07 | 0.92 | 0.92 | 0.67 |
|  | **C** | 79.17 | 79.57 | 79.37 | 0.87 | 0.89 | 0.59 | 75.27 | 79.57 | 77.42 | 0.86 | 0.86 | 0.55 |
|  | **NC** | 84.14 | 84.14 | 84.14 | 0.92 | 0.92 | 0.68 | 80.65 | 90.32 | 85.48 | **0.93** | 0.92 | 0.71 |
| **DPB**  **(SVC)** | **N** | 79.84 | 79.17 | 79.50 | 0.88 | 0.89 | 0.59 | 77.42 | 85.48 | 81.45 | 0.89 | 0.90 | 0.63 |
|  | **C** | 77.02 | 76.88 | 76.95 | 0.85 | 0.88 | 0.54 | 68.82 | 83.33 | 76.08 | 0.85 | 0.85 | 0.53 |
|  | **NC** | 82.26 | 81.86 | 82.06 | 0.90 | 0.91 | 0.64 | 79.03 | 88.17 | 83.60 | **0.91** | 0.90 | 0.68 |
| **AIB**  **(ET)** | **N** | 79.84 | 79.03 | 79.44 | 0.89 | 0.90 | 0.59 | 77.96 | 86.56 | 82.26 | 0.91 | 0.91 | 0.65 |
|  | **C** | 78.23 | 78.09 | 78.16 | 0.88 | 0.89 | 0.56 | 79.03 | 77.42 | 78.23 | 0.88 | 0.87 | 0.57 |
|  | **NC** | 82.80 | 82.12 | 82.46 | 0.91 | 0.91 | 0.65 | 80.11 | 88.17 | 84.14 | **0.93** | 0.91 | 0.69 |
| **PCB**  **(RF)** | **N** | 82.53 | 81.18 | 81.86 | 0.90 | 0.91 | 0.64 | 77.42 | 85.48 | 81.45 | 0.92 | 0.92 | 0.63 |
|  | **C** | 79.84 | 80.91 | 80.38 | 0.88 | 0.89 | 0.61 | 75.81 | 80.65 | 78.23 | 0.86 | 0.85 | 0.57 |
|  | **NC** | 82.93 | 81.45 | 82.19 | 0.91 | 0.91 | 0.64 | 82.26 | 87.63 | 84.95 | **0.93** | 0.92 | 0.70 |

**# Sn:** Sensitivity, **Sp:** Specificity, **Acc:** Accuracy, **MCC:** Matthews Correlation Coefficient, **AUC:** Area Under the Receiver Operating Characteristic curve, **AUPRC:** Area Under the Precision-Recall Curve, **AAB:** Amino acid-based binary profile, **DPB:** Dipeptide-based binary profile, **PCB:** Physico-chemical properties based binary profile, **AIB:** Amino-acid indices based binary profile, **RF:** Random Forest classifier, **ET:** Extra-tree classifier, **SVC:** Support vector classifier

**Supplementary Tabel S5: Performance of ML-models on training and validation dataset developed using four types of Binary profile-based features of N, C, NC-terminal of gram-negative ABPs**

| **Feature**  **type** | **Terminal** | **Training set** | | | | | | **Validation set** | | | | | |
| --- | --- | --- | --- | --- | --- | --- | --- | --- | --- | --- | --- | --- | --- |
|  |  | **Sn** | **Sp** | **Acc** | **AUC** | **AUPRC** | **MCC** | **Sn** | **Sp** | **Acc** | **AUC** | **AUPRC** | **MCC** |
| **AAB**  **(ET)** | **N** | 86.68 | 87.03 | 86.86 | 0.94 | 0.95 | 0.74 | 89.35 | 95.53 | 92.44 | 0.97 | 0.97 | 0.85 |
|  | **C** | 83.76 | 84.19 | 83.98 | 0.92 | 0.93 | 0.68 | 89.69 | 89.35 | 89.52 | 0.95 | 0.96 | 0.79 |
|  | **NC** | 86.77 | 87.97 | 87.37 | 0.95 | 0.95 | 0.75 | 92.44 | 94.85 | 93.64 | **0.98** | 0.98 | 0.87 |
| **DPB**  **(SVC)** | **N** | 86.25 | 85.91 | 86.08 | 0.93 | 0.94 | 0.72 | 87.97 | 91.75 | 89.86 | 0.97 | 0.97 | 0.80 |
|  | **C** | 83.42 | 83.16 | 83.29 | 0.91 | 0.93 | 0.67 | 88.32 | 88.32 | 88.32 | 0.95 | 0.96 | 0.77 |
|  | **NC** | 86.94 | 86.43 | 86.68 | 0.94 | 0.95 | 0.73 | 93.13 | 93.81 | 93.47 | **0.98** | 0.98 | 0.87 |
| **AIB**  **(ET)** | **N** | 86.86 | 86.86 | 86.86 | 0.94 | 0.95 | 0.74 | 90.03 | 94.50 | 92.27 | 0.97 | 0.97 | 0.85 |
|  | **C** | 84.02 | 84.79 | 84.41 | 0.92 | 0.93 | 0.69 | 87.97 | 87.97 | 87.97 | 0.96 | 0.96 | 0.76 |
|  | **NC** | 86.43 | 86.60 | 86.51 | 0.95 | 0.95 | 0.73 | 92.10 | 92.44 | 92.27 | **0.98** | 0.98 | 0.85 |
| **PCB**  **(ET)** | **N** | 85.83 | 86.60 | 86.21 | 0.94 | 0.94 | 0.72 | 90.72 | 93.47 | 92.10 | 0.97 | 0.97 | 0.84 |
|  | **C** | 83.25 | 83.94 | 83.59 | 0.91 | 0.92 | 0.67 | 89.00 | 85.22 | 87.11 | 0.94 | 0.95 | 0.74 |
|  | **NC** | 86.60 | 86.51 | 86.56 | 0.94 | 0.95 | 0.73 | 94.16 | 91.41 | 92.78 | **0.98** | 0.98 | 0.86 |

**# Sn:** Sensitivity, **Sp:** Specificity, **Acc:** Accuracy, **MCC:** Matthews Correlation Coefficient, **AUC:** Area Under the Receiver Operating Characteristic curve, **AUPRC:**Area Under the Precision-Recall Curve, **AAB:** Amino acid-based binary profile, **DPB:** Dipeptide-based binary profile, **PCB:** Physico-chemical properties based binary profile, **AIB:** Amino-acid indices based binary profile, **ET:** Extra-tree classifier, **SVC:** Support vector classifier

**Supplementary Tabel S6 : Performance of ML-models on training and validation dataset developed using four types of Binary profile based features of N,C,NC-terminal of gram-variable ABPs**

| **Feature type** | **Terminal** | **Training set** | | | | | | **Validation set** | | | | | |
| --- | --- | --- | --- | --- | --- | --- | --- | --- | --- | --- | --- | --- | --- |
|  |  | **Sn** | **Sp** | **Acc** | **AUC** | **AUPRC** | **MCC** | **Sn** | **Sp** | **Acc** | **AUC** | **AUPRC** | **MCC** |
| **AAB**  **(SVC)** | **N** | 85.17 | 85.16 | 85.16 | 0.92 | 0.90 | 0.70 | 83.86 | 89.26 | 86.56 | 0.93 | 0.92 | 0.73 |
|  | **C** | 82.18 | 82.15 | 82.17 | 0.89 | 0.88 | 0.64 | 79.41 | 83.03 | 81.22 | 0.89 | 0.89 | 0.63 |
|  | **NC** | 86.28 | 86.21 | 86.25 | 0.92 | 0.91 | 0.73 | 83.92 | 90.10 | 87.01 | **0.94** | 0.93 | 0.74 |
| **DPB**  **(RF)** | **N** | 45.12 | 45.28 | 45.20 | 0.48 | 0.53 | -0.10 | 64.83 | 86.14 | 75.49 | 0.86 | 0.81 | 0.52 |
|  | **C** | 80.02 | 79.35 | 79.69 | 0.88 | 0.87 | 0.59 | 76.91 | 78.69 | 77.80 | 0.86 | 0.86 | 0.56 |
|  | **NC** | 83.50 | 83.54 | 83.52 | 0.91 | 0.89 | 0.67 | 82.53 | 47.02 | 64.78 | 0.67 | 0.61 | 0.32 |
| **AIB**  **(SVC)** | **N** | 85.35 | 85.10 | 85.23 | 0.92 | 0.90 | 0.71 | 83.86 | 88.76 | 86.31 | 0.93 | 0.92 | 0.69 |
|  | **C** | 82.00 | 82.72 | 82.36 | 0.89 | 0.88 | 0.65 | 79.19 | 83.58 | 81.39 | 0.89 | 0.89 | 0.64 |
|  | **NC** | 86.42 | 86.16 | 86.29 | 0.92 | 0.91 | 0.73 | 84.53 | 90.15 | 87.34 | 0.94 | 0.93 | 0.73 |
| **PCB**  **(SVC)** | **N** | 84.71 | 84.75 | 84.73 | 0.91 | 0.90 | 0.70 | 84.09 | 88.87 | 86.48 | 0.93 | 0.92 | 0.73 |
|  | **C** | 81.27 | 80.94 | 81.11 | 0.88 | 0.87 | 0.62 | 80.52 | 82.36 | 81.44 | 0.88 | 0.89 | 0.63 |
|  | **NC** | 86.28 | 85.99 | 86.14 | 0.92 | 0.90 | 0.72 | 84.75 | 89.93 | 87.34 | 0.94 | 0.93 | 0.75 |

**# Sn:** Sensitivity, **Sp:** Specificity, **Acc:** Accuracy, **MCC:** Matthews Correlation Coefficient, **AUC:** Area Under the Receiver Operating Characteristic curve, **AUPRC:**Area Under the Precision-Recall Curve, **AAB:** Amino acid-based binary profile, **DPB:** Dipeptide-based binary profile, **PCB:** Physico-chemical properties based binary profile, **AIB:** Amino-acid indices based binary profile, **RF:** Random forest classifier, **SVC:** Support vector classifier

**Supplementary Tabel S7 : Performance of ML-models on validation dataset developed using all Binary profile-based features (AAB, AIB, PCB, DPB) of NC-terminal for GP, GN and GV ABPs**

| **Category of**  **ABPs** | **Sn** | **Sp** | **Acc** | **AUC** | **AUPRC** | **MCC** |
| --- | --- | --- | --- | --- | --- | --- |
| **Gram-positive** | 79.03 | 92.47 | 85.75 | 0.86 | 0.83 | 0.72 |
| **Gram-negative** | 92.78 | 96.22 | 94.50 | 0.95 | 0.93 | 0.89 |
| **Gram-variable** | 87.20 | 87.59 | 87.40 | 0.87 | 0.83 | 0.75 |

**# Sn:** Sensitivity, **Sp:** Specificity, **Acc:** Accuracy, **MCC:** Matthews Correlation Coefficient, **AUC:** Area Under the Receiver Operating Characteristic curve, **AUPRC:**Area Under the Precision-Recall Curve

**Supplementary Tabel S8 : Performance of ML-models on training and validation dataset developed using FastText based features for GP,GN and GV ABPs**

| **Category of**  **ABPs** | **Feature**  **(n-gram size)** | **Training set** | | | | | | **Validation set** | | | | | |
| --- | --- | --- | --- | --- | --- | --- | --- | --- | --- | --- | --- | --- | --- |
|  |  | **Sn** | **Sp** | **Acc** | **AUC** | **AUPRC** | **MCC** | **Sn** | **Sp** | **Acc** | **AUC** | **AUPRC** | **MCC** |
| **Gram-**  **positive** | **1g(ET)** | 83.20 | 83.87 | 83.54 | 0.91 | 0.91 | 0.67 | 80.65 | 87.63 | 84.14 | 0.92 | 0.91 | 0.68 |
|  | **2g(ET)** | 82.39 | 83.07 | 82.73 | 0.91 | 0.92 | 0.66 | 81.18 | 89.79 | 85.48 | 0.91 | 0.91 | 0.71 |
|  | **3g(LR)** | 78.76 | 77.96 | 78.36 | 0.87 | 0.88 | 0.57 | 72.58 | 83.33 | 77.96 | 0.84 | 0.85 | 0.56 |
|  | **2g & 3g combined**  **(ET)** | 83.07 | 82.66 | 82.86 | 0.91 | 0.92 | 0.66 | 79.57 | 85.48 | 82.53 | 0.90 | 0.90 | 0.65 |
| **Gram-**  **negative** | **1g(ET)** | 88.49 | 88.75 | 88.62 | 0.95 | 0.96 | 0.77 | 92.44 | 91.07 | 91.75 | 0.97 | 0.98 | 0.84 |
|  | **2g(ET)** | 87.29 | 87.80 | 87.54 | 0.94 | 0.95 | 0.75 | 92.10 | 90.03 | 91.07 | 0.97 | 0.98 | 0.82 |
|  | **3g(LR)** | 84.62 | 85.14 | 84.88 | 0.92 | 0.93 | 0.70 | 91.41 | 90.03 | 90.72 | 0.96 | 0.96 | 0.82 |
|  | **2g & 3g**  **combined(ET)** | 88.40 | 88.66 | 88.53 | 0.95 | 0.96 | 0.77 | 91.75 | 89.69 | 90.72 | 0.97 | 0.98 | 0.82 |
| **Gram-**  **variable** | **1g(SVC)** | 81.82 | 82.21 | 82.01 | 0.89 | 0.86 | 0.64 | 84.47 | 86.59 | 85.53 | 0.92 | 0.92 | 0.71 |
|  | **2g(SVC)** | 85.68 | 85.38 | 85.53 | 0.92 | 0.90 | 0.71 | 84.47 | 89.26 | 86.87 | 0.93 | 0.93 | 0.74 |
|  | **3g(LR)** | 83.97 | 83.92 | 83.95 | 0.91 | 0.89 | 0.68 | 81.86 | 86.14 | 84.00 | 0.92 | 0.91 | 0.68 |
|  | **2g &3g**  **combined(SVC)** | 86.63 | 86.24 | 86.44 | 0.92 | 0.91 | 0.73 | 83.92 | 89.20 | 86.56 | 0.93 | 0.92 | 0.73 |

**# Sn:** Sensitivity, **Sp:** Specificity, **Acc:** Accuracy, **MCC:** Matthews Correlation Coefficient, **AUC:** Area Under the Receiver Operating Characteristic curve, **AUPRC:**Area Under the Precision-Recall Curve

**Supplementary Tabel S9 : Performance of hybrid model(BLAST+ML) on the validation set of gram-positive ABPs independent at different e-value**

| **E-value** | **Correct hits**  **(BLAST+ML)** | **Sn** | **Sp** | **Acc** | **AUC** | **AUPRC** | **MCC** |
| --- | --- | --- | --- | --- | --- | --- | --- |
| **1.00E-20** | **86.02%** | **81.72** | **90.32** | **86.02** | **0.94** | **0.93** | **0.72** |
| **1.00E-10** | 85.75% | 81.18 | 90.32 | 85.75 | 0.93 | 0.92 | 0.72 |
| **1.00E-06** | 86.29% | 82.26 | 90.32 | 86.29 | 0.93 | 0.91 | 0.73 |
| **0.0001** | 86.83% | 83.33 | 90.32 | 86.83 | 0.93 | 0.91 | 0.74 |
| **0.001** | 87.90% | 85.48 | 90.32 | 87.90 | 0.93 | 0.91 | 0.76 |
| **0.01** | 88.44% | 86.02 | 90.86 | 88.44 | 0.93 | 0.92 | 0.77 |
| **0.1** | 87.90% | 86.56 | 89.25 | 87.90 | 0.93 | 0.92 | 0.76 |
| **1** | 86.56% | 87.10 | 86.02 | 86.56 | 0.93 | 0.92 | 0.73 |
| **10** | 81.18% | 83.87 | 78.49 | 81.18 | 0.90 | 0.89 | 0.62 |
| **50** | 80.11% | 83.33 | 76.88 | 80.11 | 0.90 | 0.89 | 0.60 |
| **100** | 80.38% | 83.33 | 77.42 | 80.38 | 0.90 | 0.89 | 0.61 |
| **200** | 80.11% | 83.33 | 76.88 | 80.11 | 0.90 | 0.89 | 0.60 |
| **1000** | 80.38% | 83.87 | 76.88 | 80.38 | 0.91 | 0.89 | 0.61 |
| **Only ML** | 85.48% | 80.65 | 90.32 | 85.48 | 0.93 | 0.92 | 0.71 |

**# Sn:** Sensitivity, **Sp:** Specificity, **Acc:** Accuracy, **MCC:** Matthews Correlation Coefficient, **AUC:** Area Under the Receiver Operating Characteristic curve, **AUPRC:**Area Under the Precision-Recall Curve

**Supplementary Tabel S10 : Performance of hybrid model (BLAST+ML) on the validation set of gram-negative ABPs independent at different e-value**

| **E-value** | **Correct hits**  **(BLAST+ML)** | **Sn** | **Sp** | **Acc** | **AUC** | **AUPRC** | **MCC** |
| --- | --- | --- | --- | --- | --- | --- | --- |
| **1.00E-20** | 93.30% | 92.44 | 94.16 | 93.30 | 0.98 | 0.98 | 0.87 |
| **1.00E-10** | 93.99% | 94.16 | 93.81 | 93.99 | 0.98 | 0.97 | 0.88 |
| **1.00E-06** | 94.16% | 94.50 | 93.81 | 94.16 | 0.97 | 0.97 | 0.88 |
| **0.0001** | 94.16% | 94.16 | 94.16 | 94.16 | 0.97 | 0.97 | 0.88 |
| **0.001** | 94.33% | 94.50 | 94.16 | 94.33 | 0.97 | 0.97 | 0.89 |
| **0.01** | 94.67% | 94.85 | 94.50 | 94.67 | 0.97 | 0.97 | 0.89 |
| **0.1** | 94.85% | 94.85 | 94.85 | 94.85 | 0.97 | 0.97 | 0.90 |
| **1** | 94.16% | 93.81 | 94.50 | 94.16 | 0.97 | 0.97 | 0.88 |
| **10** | 91.41% | 92.10 | 90.72 | 91.41 | 0.96 | 0.96 | 0.83 |
| **50** | 90.21% | 90.72 | 89.69 | 90.21 | 0.96 | 0.96 | 0.80 |
| **100** | 90.03% | 90.38 | 89.69 | 90.03 | 0.96 | 0.96 | 0.80 |
| **200** | 89.86% | 90.03 | 89.69 | 89.86 | 0.96 | 0.96 | 0.80 |
| **1000** | 89.18% | 88.66 | 89.69 | 89.18 | 0.96 | 0.96 | 0.78 |
| **Only ML** | 93.64% | 92.44 | 94.85 | 93.64 | 0.98 | 0.98 | 0.87 |

**# Sn:** Sensitivity, **Sp:** Specificity, **Acc:** Accuracy, **MCC:** Matthews Correlation Coefficient, **AUC:** Area Under the Receiver Operating Characteristic curve, **AUPRC:**Area Under the Precision-Recall Curve

**Supplementary Tabel S11 : Performance of hybrid model (BLAST+ML) on the validation set of gram-variable ABPs independent at different e-value**

| **E-value** | **Correct hits**  **(BLAST+ML)** | **Sn** | **Sp** | **Acc** | **AUC** | **AUPRC** | **MCC** |
| --- | --- | --- | --- | --- | --- | --- | --- |
| **1.00E-20** | 87.26% | 86.53 | 88.59 | 87.56 | 0.93 | 0.92 | 0.75 |
| **1.00E-10** | 87.37% | 87.15 | 88.31 | 87.73 | 0.92 | 0.87 | 0.75 |
| **1.00E-06** | 87.59% | 86.76 | 88.93 | 87.84 | 0.92 | 0.86 | 0.76 |
| **0.0001** | 87.62% | 86.81 | 88.76 | 87.79 | 0.92 | 0.86 | 0.76 |
| **0.001** | 87.62% | 86.59 | 88.81 | 87.70 | 0.92 | 0.87 | 0.75 |
| **0.01** | 87.40% | 86.09 | 88.87 | 87.48 | 0.92 | 0.87 | 0.75 |
| **0.1** | 87.34% | 85.87 | 88.76 | 87.31 | 0.92 | 0.88 | 0.75 |
| **1** | 86.84% | 85.14 | 88.26 | 86.70 | 0.92 | 0.88 | 0.73 |
| **10** | 83.81% | 81.36 | 85.03 | 83.19 | 0.91 | 0.88 | 0.66 |
| **50** | 82.39% | 79.19 | 84.14 | 81.66 | 0.91 | 0.88 | 0.63 |
| **100** | 82.03% | 78.91 | 83.75 | 81.33 | 0.91 | 0.88 | 0.63 |
| **200** | 81.91% | 78.80 | 83.64 | 81.22 | 0.91 | 0.88 | 0.63 |
| **1000** | 81.80% | 78.58 | 83.58 | 81.08 | 0.91 | 0.88 | 0.62 |
| **Only ML** | 87.01% | 83.92 | 90.10 | 87.01 | 0.94 | 0.93 | 0.74 |

**# Sn:** Sensitivity, **Sp:** Specificity, **Acc:** Accuracy, **MCC:** Matthews Correlation Coefficient, **AUC:** Area Under the Receiver Operating Characteristic curve, **AUPRC:**Area Under the Precision-Recall Curve

**Supplementary Tabel S12 : Performance of hybrid model(MOTIF+ML) on the validation set of gram-positive ABPs independent set at different frequencies**

| **Frequency of a motif** | **Correct hits**  **(MOTIF+ML)** | **Sn** | **Sp** | **Acc** | **AUC** | **AUPRC** | **MCC** |
| --- | --- | --- | --- | --- | --- | --- | --- |
| **fp10_motif** | 84.68% | 83.33 | 86.02 | 84.68 | 0.91 | 0.88 | 0.69 |
| **fp20_motif** | 85.22% | 81.72 | 88.71 | 85.22 | 0.92 | 0.90 | 0.71 |
| **fp30_motif** | 85.22% | 81.18 | 89.25 | 85.22 | 0.92 | 0.90 | 0.71 |
| **Only ML** | **85.48%** | **80.65** | **90.32** | **85.48** | **0.93** | **0.92** | **0.71** |

**# Sn:** Sensitivity, **Sp:** Specificity, **Acc:** Accuracy, **MCC:** Matthews Correlation Coefficient, **AUC:** Area Under the Receiver Operating Characteristic curve, **AUPRC:**Area Under the Precision-Recall Curve

**Supplementary Tabel S13 : Performance of hybrid model(MOTIF+ML) on the validation set of gram-negative ABPs independent set at different frequencies**

| **Frequency of a motif** | **Correct hits**  **(MOTIF+ML)** | **Sn** | **Sp** | **Acc** | **AUC** | **AUPRC** | **MCC** |
| --- | --- | --- | --- | --- | --- | --- | --- |
| **fp10_motif** | 90.21% | 93.47 | 86.94 | 90.21 | 0.96 | 0.95 | 0.81 |
| **fp20_motif** | 92.27% | 92.78 | 91.75 | 92.27 | 0.97 | 0.97 | 0.85 |
| **fp30_motif** | 93.30% | 92.78 | 93.81 | 93.30 | 0.97 | 0.97 | 0.87 |
| **Only ML** | **93.64%** | **92.44** | **94.85** | **93.64** | **0.98** | **0.98** | **0.86** |

**# Sn:** Sensitivity, **Sp:** Specificity, **Acc:** Accuracy, **MCC:** Matthews Correlation Coefficient, **AUC:** Area Under the Receiver Operating Characteristic curve, **AUPRC:**Area Under the Precision-Recall Curve

**Supplementary Tabel S14 : Performance of hybrid model(MOTIF+ML) on the validation set of gram-variable ABPs independent set at different frequencies**

| **Frequency of a motif** | **Correct hits**  **(MOTIF+ML)** | **Sn** | **Sp** | **Acc** | **AUC** | **AUPRC** | **MCC** |
| --- | --- | --- | --- | --- | --- | --- | --- |
| **fp10_motif** | 86.03% | 86.70 | 85.36 | 86.03 | 0.93 | 0.92 | 0.72 |
| **fp20_motif** | 87.01% | 85.75 | 88.26 | 87.01 | 0.94 | 0.94 | 0.74 |
| **fp30_motif** | 87.28% | 85.48 | 89.09 | 87.28 | 0.94 | 0.93 | 0.75 |
| **Only ML** | **87.01%** | **83.92** | **90.10** | **87.01** | **0.94** | **0.93** | **0.74** |

**# Sn:** Sensitivity, **Sp:** Specificity, **Acc:** Accuracy, **MCC:** Matthews Correlation Coefficient, **AUC:** Area Under the Receiver Operating Characteristic curve, **AUPRC:**Area Under the Precision-Recall Curve
